# Classifying *Drosophila* Olfactory Projection Neuron Subtypes by Single-cell RNA Sequencing

**DOI:** 10.1101/145045

**Authors:** Hongjie Li, Felix Horns, Bing Wu, Qijing Xie, Jiefu Li, Tongchao Li, David Luginbuhl, Stephen R. Quake, Liqun Luo

**Author notes:** These authors contributed equally to the work (Hongjie Li, Felix Horns). Corresponding authors (L.L.); (S.R.Q.).

## Abstract

How a neuronal cell type is defined and how this relates to its transcriptome are still open questions. The *Drosophila* olfactory projection neurons (PNs) are among the best-characterized neuronal types: Different PN classes target dendrites to distinct olfactory glomeruli and PNs of the same class exhibit indistinguishable anatomical and physiological properties. Using single-cell RNA-sequencing, we comprehensively characterized the transcriptomes of 40 PN classes and unequivocally identified transcriptomes for 6 classes. We found a new lineage-specific transcription factor that instructs PN dendrite targeting. Transcriptomes of closely-related PN classes exhibit the largest difference during circuit assembly, but become indistinguishable in adults, suggesting that neuronal subtype diversity peaks during development. Genes encoding transcription factors and cell-surface molecules are the most differentially expressed, indicating their central roles in specifying neuronal identity. Finally, we show that PNs use highly redundant combinatorial molecular codes to distinguish subtypes, enabling robust specification of cell identity and circuit assembly.

## Introduction

The nervous system comprises many types of neurons with varied locations, input and output connections, neurotransmitter phenotypes, intrinsic properties, and physiological and behavioral functions. Recent transcriptome analyses, especially from single cells, have provided important criteria to define a cell type. Indeed, single-cell RNA-sequencing (RNA-seq) has been used to classify neurons in various parts of the mammalian nervous system (e.g., Darmanis et al., 2015; Johnson et al., 2015; Usoskin et al., 2015; Zeisel et al., 2015; Foldy et al., 2016; Fuzik et al., 2016; Shekhar et al., 2016; Tasic et al., 2016), but the extent to which it is useful to define subtypes of neurons and the relationship between cell type and neural connectivity is unclear in most cases. Indeed, what constitutes a neuronal type in many parts of the nervous system is still an open question (Johnson and Walsh, 2017).

The *Drosophila* olfactory circuit offers an excellent system to investigate the relationship between transcriptomes and neuronal cell types. 50 classes of olfactory receptor neurons (ORNs) form one-to-one connections with 50 classes of second-order projection neurons (PNs) in the antennal lobe in 50 discrete glomeruli, forming 50 parallel information processing channels (Figure 1A; Vosshall and Stocker, 2007; Su et al., 2009; Wilson, 2013). Each ORN class is defined by expression of 1–2 unique olfactory receptor gene(s) and by the glomerulus to which their axons converge (Gao et al., 2000; Vosshall et al., 2000; Couto et al., 2005; Silbering et al., 2011). Correspondingly, each PN class is also defined by the glomerulus within which their dendrites elaborate (Stocker et al., 1990; Jefferis et al., 2001), which correlates strongly with the axonal arborization patterns at the lateral horn, a higher olfactory center (Marin et al., 2002; Wong et al., 2002; Jefferis et al., 2007). Furthermore, while on average ~60 ORNs and ~3 PNs form many hundreds of synapses within a single glomerulus (Mosca and Luo, 2014), every ORN forms synapses with every PN to convey the same type of olfactory information (Kazama and Wilson, 2009; Tobin et al., 2017). Indeed, PNs that project to the same glomerulus exhibit indistinguishable electrophysiological properties and olfactory responses (Kazama and Wilson, 2009). Thus, one can define each class of PN as a specific neuronal type (or subtype, if all PNs are collectively considered a cell type) with confidence that each class has unique connectivity, physiological response properties, and function, whereas individual PNs within the same class most likely do not differ. In other words, the ground truth of cell types for fly PNs is one of the best defined in the nervous system.

**Figure 1.**
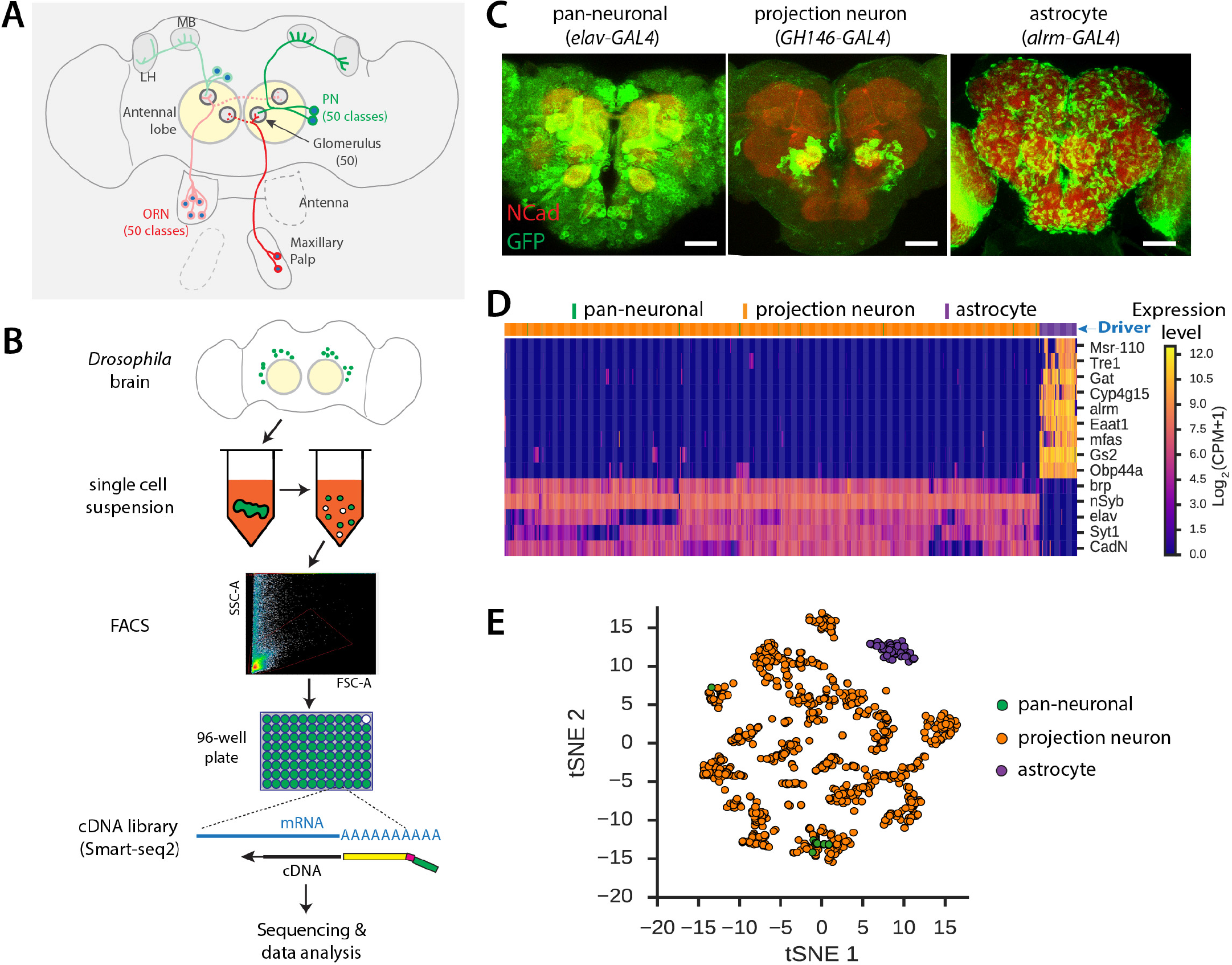
Single-cell RNA-seq Protocol for the *Drosophila* Pupal Brain. (A) Schematic of the fly olfactory system organization. Olfactory receptor neurons (ORNs), whose cell bodies are located in the antenna and maxillary palp, expressing the same receptor (same color) target their axons to the same glomerulus in the antennal lobe. Projection neuron (PN) dendrites also target to single glomeruli, and their axons project to the mushroom body (MB) and lateral horn (LH). (B) Schematic of single cell RNA-seq protocol for *Drosophila* brain cells. See text for details. (C) Representative confocal images of *Drosophila* central brains that are labeled by *UAS-mCD8GFP* crossed with pan-neuronal driver *elav-GAL4* (24h APF), olfactory projection neuron driver *GH146-GAL4* (24h APF), and astrocyte driver *alrm-GAL4* (72h APF). N-cadherin (Ncad) staining was used to label neuropil (red). Scale bar, 50 µm. (D) Heat map showing expression levels of genes that are specific for neurons or astrocytes. Each column is an individual cell. 12 *elav-GAL4*+, 67 *alrm-GAL4*+, and 946 *GH146-GAL4*+ cells are shown, and the driver used to label the cell is indicated by the color above the column. Cell type-specific genes are enriched in astrocytes (top 9) and neurons (bottom 5). Expression levels are indicated by the color bar (CPM, counts per million). Cells and genes were ordered using hierarchical clustering. (E) Visualization of astrocyte and neuron populations using t-distributed Stochastic Neighbor Embedding (tSNE). Each dot is one cell. Color indicates the driver used to label the cell. See also Figure S1.

While single-cell RNA-seq has been increasingly applied to study mammalian neurons (Johnson and Walsh, 2017), to our knowledge it has not been systematically applied to the *Drosophila* nervous system. This is likely due to the small size of fly cell bodies (3–4 µm diameter for a typical neuron), making it difficult to obtain enough mRNA from single cells for reliable cDNA library preparation (Crocker et al., 2016). Recent technical advances have significantly increased cDNA library yield from small amounts of RNA (Goetz and Trimarchi, 2012; Picelli et al., 2014), providing an opportunity to interrogate *Drosophila* olfactory neurons using single-cell RNA-seq. Here, we have established a robust single-cell RNA-seq protocol for neurons and glial cells in the *Drosophila* brain. We sequenced 1,277 single cells of 40 classes of PNs, focusing mostly on a developmental stage when PNs are establishing wiring specificity. Using an unsupervised machine-learning algorithm that we developed, we could assign these PNs to 30 distinct clusters. Based on known and newly identified specific markers, we unambiguously mapped 6 PN classes derived from two neuroblast lineages to specific clusters. We also identified new lineage-specific transcription factors, finding that the homeodomain-containing C15 instructs lineage-specific dendrite targeting. By comparing transcriptomes of identified PN classes across developmental stages, we found that two closely-related PN classes have distinct transcriptome signatures at stages when they are establishing wiring specificity, but their transcriptomes become indistinguishable in adults. Systematic analyses revealed that PN ensembles employ a combinatorial, redundant code to distinguish different subtypes, and that genes encoding transcription factors and cell-surface molecules are among the most differentially expressed in different PN subtypes and are highly informative for distinguishing subtypes.

## Results

### A Robust Single-cell RNA-seq Protocol for the *Drosophila* Pupal Brain

We developed an RNA-seq protocol for single cells dissociated from the *Drosophila* pupal brain (Figure 1B; Experimental Procedures). Briefly, pupal brains containing cells labeled by mCD8GFP driven from specific GAL4 lines were manually dissected, and single-cell suspensions were prepared following a method modified after Tan et al. (2015). We confirmed using phase-contrast microscopy that nearly all individual cells were separated (Figure S1A). Single GFP-positive cells were sorted into individual wells of 96-well plates using Fluorescence Activated Cell Sorting (FACS). We reverse transcribed poly(A)-RNA and amplified full-length cDNA using a modified SMART-seq2 protocol with an increased number of PCR cycles to offset the small amount of mRNA in each cell (Picelli et al., 2014). We prepared sequencing libraries from cDNA using tagmentation (Nextera XT) and sequenced them using the Illumina platform.

In pilot experiments, we sequenced a small number of cells from *Drosophila* pupal brains that were labeled by the pan-neuronal driver *elav-GAL4* (Lin and Goodman, 1994) or by the astrocyte driver *alrm-GAL4* (Doherty et al., 2009) (Figure 1C). We analyzed them together with a much larger number of olfactory PNs labeled by the *GH146-GAL4* driver, which is expressed in 40 of 50 PN classes (Stocker et al., 1997; Jefferis et al., 2001). Approximately 5% of GFP-labeled cells within the brain could be recovered as single cells sorted into plates after brain dissociation and FACS, and 90% of neurons yielded high-quality cDNA after reverse transcription (Figure S1B and S1D). Cells from all three drivers were sequenced to a depth of ~1 million reads per cell and 1000–4000 genes were detected in each cell (Figure S1C). To evaluate the quality of RNA-seq data, we examined expression of 5 known neuronal markers (*brp, nSyb, elav, Syt1, and CadN*) and 4 known astrocyte markers (*alrm, Eaat1, Gat, and Gs2*) (Doherty et al., 2009; Sinakevitch et al., 2010; Stork et al., 2014), and found that they were specifically expressed in the corresponding cell types (Figure 1D). In addition, we identified 5 new genes (*Msr-110, tre1, Cyp4g15, mfas, Obp44a*) that were expressed in pupal astrocytes but not in neurons (Figure 1D). Unbiased clustering based on transcriptome profiles readily distinguished neurons and astrocytes (Figure 1E). These data support the reliability of our single-cell RNA-seq protocol for analyzing cell types and transcriptomes in *Drosophila* pupal brain (Figure S1D), and suggest that it would be applicable to other tissues and developmental stages.

### Single-cell RNA-seq Analysis of *GH146-GAL4*+ Projection Neurons (PNs)

We previously showed that *GH146-GAL4*+ (*GH146*+ hereafter) PNs are derived from three separate neuroblast lineages whose cell bodies are located anterodorsal, lateral, or ventral to the antennal lobe neuropil (Figure 2A; Jefferis et al., 2001). The anterodorsal and lateral lineages give rise to uniglomerular excitatory PNs (adPNs and lPNs) that target to complementary glomeruli (Jefferis et al., 2001), whereas the ventral lineage produces GABAergic inhibitory PNs (vPNs) (Liang et al., 2013). We sequenced 1046 single *GH146*+ PNs at 24–30 hours after puparium formation (h APF). At this stage, PNs are refining their dendrite targeting in the antennal lobe; their dendrites further serve as targets for ORN axons that will invade the antennal lobe and establish one-to-one connections in the next 24 hours (Jefferis et al., 2004). 946 cells passed the filter of expressing at least 4 out of 6 markers (*mCD8GFP* and 5 neuronal markers in Figure 1D). Housekeeping genes (e.g., *Act5C* and *Rp49*) were reliably detected in all cells. About 50% of cells co-expressed male-specific genes, *RNA on the X 1* and *RNA on the X 2* (Meller et al., 1997) (Figure S2A), as expected given that we did not discriminate sex.

**Figure 2.**
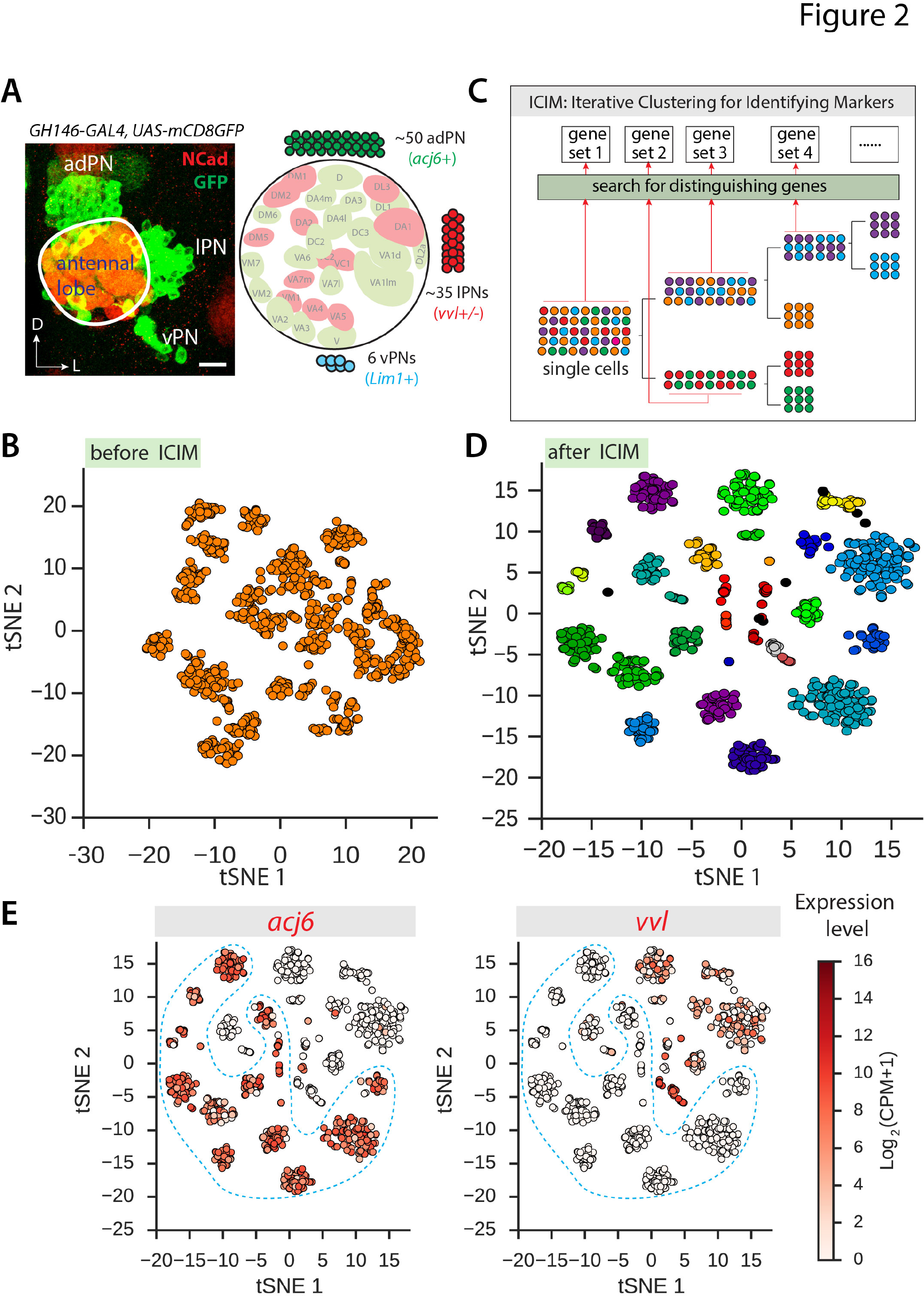
Single-cell RNA-seq Analysis for *GH146*+ Projection Neurons. (A) Representative image (confocal projection) and schematic of *GH146-GAL4*+ olfactory projection neurons (PNs), which include (per antennal lobe) 50 adPNs *(acj6+)*, 35 lPNs (*vvl*+; expression of *vvl* begins to decrease in lPNs from 18h APF) and 6 vPNs (Lim1+). The cell bodies of adPNs, lPNs, and vPNs are located respectively anterodorsal, lateral, and ventral to the antennal lobe neuropil (circled), which was labeled by N-cadherin (Ncad) staining in red. All *GH146*+ adPNs and lPNs send dendrites to a single, specific glomerulus. The schematic shows the stereotyped locations of a large subset of glomeruli (named according to their locations; Laissue et al., 1999) in a 2D projection, color-coded according to whether they are targeted by adPNs or lPNs. Scale bar, 20 µm. D, dorsal; L, lateral. (B) Visualization of *GH146*+ PN cells using dimensional reduction by PCA followed by tSNE. Each dot is one cell. Cells are arranged according to transcriptome similarity. (C) Schematic of Iterative Clustering for Identifying Markers (ICIM), an unsupervised machine-learning algorithm for identifying genes that distinguish cell types. See text for details. (D) Visualization of *GH146*+ PN cells using tSNE based on 561 genes identified using ICIM. Each dot is one cell. Cells are arranged according to similarity of expression profiles of the selected genes. Cells are colored by cluster identity, as determined using HDBSCAN. *GH146*+ adPNs and lPNs form 30 distinct clusters. (E) Visualization of *GH146*+ PN cells arranged using ICIM and tSNE as in Figure 2D and colored according to *acj6* and *vvl* expression level (CPM, counts per million). *acj6* and *vvl* (encoding lineage-specific transcription factors for adPNs and lPNs, respectively) are expressed in *GH146*+ PNs in a mutually exclusive manner. ~60% of *GH146-GAL4*+ cells are *acj6*+ (see Figure 2A), consistent with previous findings (Komiyama et al., 2003). See also Figure S2.

We tried to distinguish PN subtypes using conventional dimensional reduction and clustering methods based on Principal Component Analysis (PCA) and t-distributed Stochastic Neighbor Embedding (tSNE) (van der Maaten and Hinton, 2008). This approach identified only ~12 distinct clusters of PNs (Figure 2B). The inability to resolve more distinct clusters is likely due to the limited sensitivity of these analysis methods to distinguish cell types having highly similar transcriptomes, which is likely the case for different PN classes. To address this challenge, we developed an unsupervised machine-learning algorithm, which we call Iterative Clustering for Identifying Markers (ICIM), to identify genes that distinguish PN classes (see Experimental Procedures). Briefly, this algorithm searches for genes having the highest expression variability within a cell population, partitions the cells into two subpopulations using clustering based on these genes, then iteratively repeats the search on each subpopulation. Iteration continues until distinct subpopulations cannot be separated because gene expression patterns within the population are homogeneous (Figure 2C). Stopping criteria are defined in an unbiased manner without supervision. Genes identified using ICIM were then used for dimensionality reduction using tSNE and clustering using HDBSCAN, a hierarchical density-based clustering algorithm (Campello et al., 2013). Applying ICIM to the transcriptomes of *GH146*+ PNs, we identified 561 genes that segregate the 946 *GH146*+ cells into 33 distinct clusters (Figure S2B and S2C).

Based on known markers, we assigned cell type identities to three clusters. Among the 33 clusters, two clusters comprising 11 cells expressed known markers for vPNs [*Gad1*+, which encodes a GABA biosynthetic enzyme, and *Lim1*+, which encodes a transcription factor that is expressed in vPNs but not adPNs or lPNs (Komiyama and Luo, 2007)] (Figure S2B). Besides PNs, *GH146-GAL4* also labels the anterior paired lateral (APL) neurons in the central brain (Figure S2D). A third cluster, comprising 33 neurons, expressed *VGlut* robustly (Figure S2C). We observed that *VGlut-GAL4* (after intersecting with *GH146-Flp*; see below for details of this intersectional strategy) specifically labeled APL neurons but not PNs (Figure S2E), suggesting that this *VGlut*+ population consists of APL neurons. Because we were interested primarily in *GH146*+ adPNs and lPNs, we removed vPNs and APL neurons from subsequent analysis.

We focused our analysis on the 902 identified *GH146*+ adPNs and lPNs, each of which targets dendrites to a single glomerulus (Jefferis et al., 2001). Collectively, *GH146*+ adPNs and lPNs target 40 glomeruli. Clustering analysis using ICIM and tSNE identified 30 distinct clusters (Figure 2D). The number of cells belonging to each cluster varied from 5 to 108, partly reflecting the fact that different PN classes contain different cell numbers ranging from 1 to 7 per antennal lobe. Among the 40 *GH146*+ PN classes, 17 have only a single cell per antennal lobe (Yu et al., 2010; Lin et al., 2012). It is likely that we did not sample a sufficient number of cells to separate these rare PN classes into distinct clusters, contributing to our inability to identify 40 clusters corresponding to the 40 PN classes.

We previously showed that two transcription factors, Abnormal chemosensory jump 6 (Acj6) and Ventral veins lacking (Vvl; also known as Drifter), are expressed in adPNs and lPNs, respectively (Figure 2A), and instruct lineage-specific dendrite targeting (Komiyama et al., 2003). The expression of Acj6 in adPNs persists in all pupal and adult stages, while the expression of Vvl can be detected in all lPNs in the early pupa but begins to decrease from 18h APF (Komiyama et al., 2003). Indeed, our single-cell RNA-seq analysis revealed that *acj6* and *vvl* were expressed in a mutually exclusive manner (Figure 2E and S2F). Among the 30 clusters of adPNs and lPNs, 18 clusters (60%) expressed *acj6* but not *vvl*, and thus represent adPNs (Figure 2E and S2F). The remaining 12 clusters expressed either *vvl* alone, or neither *acj6* nor *vvl* (Figure 2E and S2F), and were likely lPNs since the Vvl protein is specifically expressed in lPNs and its expression declines after 18h APF.

In summary, single-cell RNA-seq revealed 30 distinct clusters of *GH146*+ adPNs and lPNs expressing lineage markers in a manner consistent with previous knowledge. Transcription factors, which at the protein level are thought to be generally present in low-abundance (Ghaemmaghami et al., 2003), were reliably detected within these cells and could be used to assign lineage identity, supporting the specificity and sensitivity of our method.

### Matching Clusters to PN Classes Using Known Markers

Next we attempted to map the correspondence between transcriptome-based PN clusters and glomerular-target-based PN classes by leveraging drivers that label specific PN classes. In principle, we could use a driver that labels one specific PN class among the *GH146*+ PNs to conduct single-cell RNA-seq and map the cells into the clusters identified among the *GH146*+ PNs. If the cells labeled by the driver could be unambiguously mapped into specific *GH146*+ PN cluster(s), we could unequivocally assign the PN class identity to the cluster(s). Drivers used for this strategy must be turned on early enough, allowing cell collection at 24–30h APF, the same stage as the sequenced *GH146*+ PNs. Drivers must also be expressed later (>48h APF) when PN wiring is largely complete and glomeruli can be identified based on their stereotyped size, shape, and location (Jefferis et al., 2004).

We tested this strategy first using *91G04-GAL4* from the Janelia GAL4 collection (Jenett et al., 2012); we found this driver to be robustly expressed in PNs at 24h APF from a GAL4 screen. To limit expression only to PNs, we utilized an intersectional strategy by combining *91G04-GAL4* with *GH146-Flp* (Potter et al., 2010) and *UAS-FRT-STOP-FRT-mCD8GFP* (Hong and Luo, 2009), such that only cells that express both *91G04-GAL4* and *GH146-Flp* will express mCD8GFP (we refer to this strategy as intersecting with *GH146-Flp* hereafter). This intersection resulted in expression of mCD8GFP in just two adPNs per brain hemisphere, both of which project dendrites to the DC2 glomerulus (Figure 3A). We sequenced 23 *91G04*+ PNs at 24–30h APF and performed clustering analysis using ICIM and tSNE together with the *GH146*+ cells. We found that all *91G04*+ PNs mapped to one *GH146*+ cluster (Figure 3C; Cluster #1). All of the *91G04*+ cells could also be unambiguously mapped to this *GH146*+ cluster using a random forest classifier (data not shown). We conclude that Cluster #1 corresponds to DC2 PNs.

**Figure 3.**
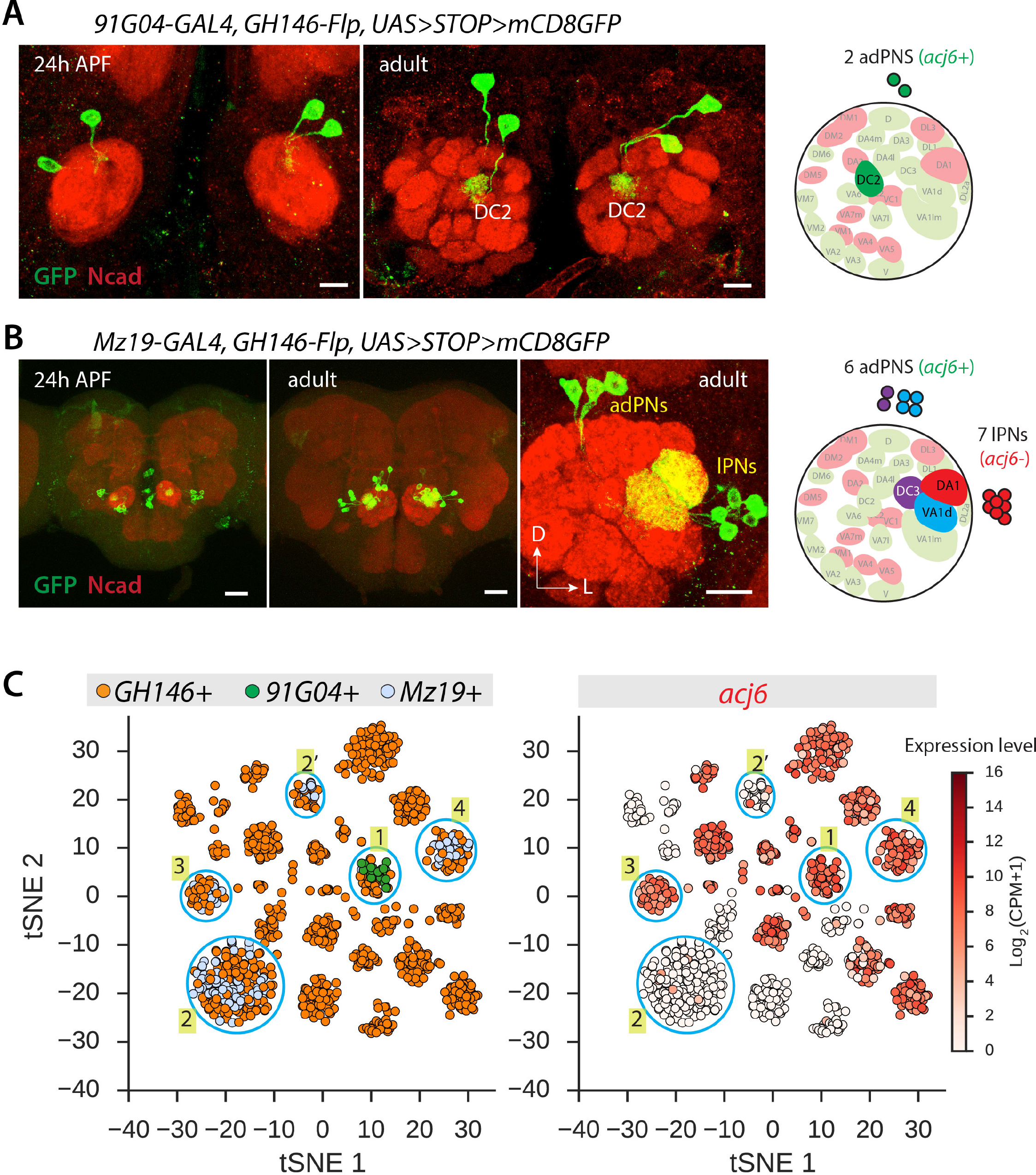
Assign Clusters to PN Classes Using Known Markers. (A and B) Intersecting *GH146-Flp* with *91G04-GAL4* (A) or with *Mz19-GAL4* (B) labels only one PN class (DC2) or 3 PN classes (VA1d and DC3 that are *acj6*+ adPNs; DA1 that are *acj6−* lPNs), respectively. Both intersections label similar numbers of PNs at 24h APF and in adults. N-cadherin (Ncad) staining was used to label neuropil (red). Schematics of these two drivers are shown on the right. Scale bar for (A), 20 µm; for (B), 50 µm, 50 µm and 20 µm. (C) Visualization of *GH146+, 91G04*+, and *Mz19*+ PN cells using tSNE based on 561 genes previously identified using ICIM (Figure 2D). Each dot is one cell. Cells are arranged according to similarity of expression profiles of the selected genes. In the left panel, cells are colored according to driver. In the right panel, cells are colored by expression level of *acj6* (CPM, counts per million). We mapped Cluster #1 to DC2 PNs, corresponding to *91G04*+ cells. DA1 PNs *(Mz19+/acj6−)* map to two clusters, #2 and #2’. VA1d and DC3 PNs *(Mz19+/acj6+)* map to two clusters, #3 and #4; we elucidated which glomerulus corresponds to which cluster in Figure 4.

We next focused on a second driver, *Mz19-GAL4* (Jefferis et al., 2004), which is expressed from 24h APF to adulthood (Figure 3B). After intersecting with *GH146-Flp, Mz19-GAL4* labels three classes of PNs: adPNs that project to VA1d and DC3 (*acj6*+), and lPNs that project to DA1 *(acj6−)*. We sequenced 123 *Mz19*+ cells at 24–30h APF, and performed clustering analysis together with *GH146*+ cells. We found that the *Mz19*+ cells mapped to four clusters of *GH146*+ cells (Figure 3C). The Mz19+/acj6− cells, corresponding to DA1 PNs, mapped to two clusters (#2 and #2’). One of these clusters (#2) contained substantially more cells than the other (#2’). This finding suggests that both clusters correspond to DA1 PNs, a notion that we explore further in the next section. The *Mz19+/acj6*+ cells, corresponding to VA1d and DC3 PNs, mapped to two clusters of *GH146*+ cells (#3 and #4) (Figure 3C). We conclude that Clusters #3 and #4 correspond to VA1d and DC3 PNs. In the next section, we present results that establish a one-to-one correspondence between these clusters and PN classes using further experiments. Together, these results demonstrate that single-cell RNA-seq and existing genetic drivers can be harnessed to rapidly map PN classes to transcriptome identities.

### Matching Clusters to PN Classes Using Newly Identified Markers

To map additional PN transcriptome clusters to glomerular classes, we asked whether we could identify new markers and genetic drivers based on single-cell transcriptome data. While searching for marker genes which were predominantly expressed in a single cluster, we identified *terribly reduced optic lobes (trol;* Figure 4A). We examined reporter expression driven by a *trol* driver in PNs by intersecting an existing *trol-GAL4 (NP5103-GAL4*, inserted into an intron of *trol*) with *GH146-Flp*. We found that *trol-GAL4* was expressed in 2–3 adPNs at both 24h APF and 72h APF, and these PNs all sent dendrites to the VM2 glomerulus at 72 h APF (Figure 4B). We hypothesized that if *trol-GAL4* recapitulated the endogenous expression of *trol*, we could then assign the *trol*+ cluster to VM2 PNs. To verify this, we sequenced 28 *trol-GAL4*+ cells (after intersecting with *GH146-Flp*). We found that these cells indeed mapped to the original *trol*+ Cluster #5 (Figure 4C). These data indicate that *trol-GAL4* mimics endogenous *trol* expression and that the Cluster #5 corresponds to VM2 PNs.

**Figure 4.**
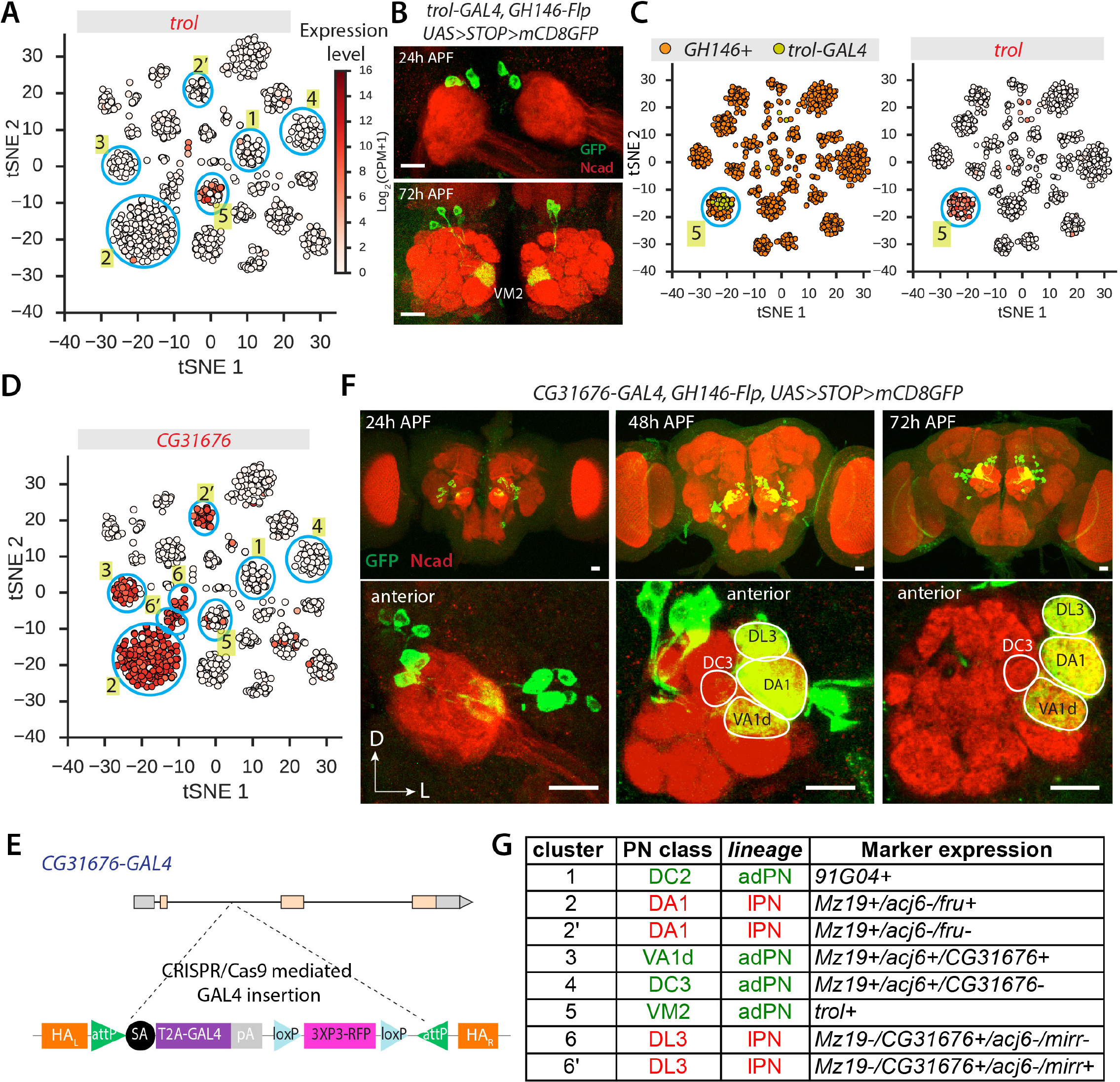
Assign Clusters to PN Classes Using Newly Identified Markers. (A) Visualization of *GH146*+ PN cells using tSNE based on 561 genes previously identified using ICIM (Figure 2D) showing expression of *trol* (CPM, counts per million) enriched in one cluster (#5). Clusters #1–4 from Figure 3 are also indicated. (B) After intersecting with *GH146-Flp, trol-GAL4* labels 2–3 PNs in each hemisphere at both 24h and 72h APF. These PNs all project dendrites to the VM2 glomerulus as shown at 72h APF. Scale bar, 20 µm. (C) Visualization of *GH146*+ and *trol*+ PN cells using tSNE based on 561 genes previously identified using ICIM (Figure 2D). In the left panel, cells are colored according to driver. In the right panel, cells are colored by expression level of *trol* (see color bar in Figure 4A; CPM, counts per million). *trol-GAL4*+ PNs map to one *GH146*+ PN cluster (left), which displays high expression of the endogenous gene *trol* (right). This cluster (#5) is mapped to VM2 PNs. Note that because of the stochastic nature of tSNE the arrangement of cells and clusters changes when a new group of cells is added (compare cluster #5 on Figure 3A and 3C). (D) Visualization of *GH146*+ PN cells using tSNE as in Figure 4A showing expression of *CG31676* (CPM, counts per million; see color bar in Figure 4A). *CG31676* is highly expressed in one of two *Mz19+/acj6*+ clusters (VA1d and DC3, clusters #3 and #4). Several other clusters also express *CG31676*, including the DA1 clusters (#2 and #2’). Clusters #6 and #6’ also express for *CG31676*, but are *acj6−* (see Figure S3B). (E) Schematic of CRISPR/Cas9 mediated insertion of *T2A-GAL4* into the first intron of *CG31676*. (F) *CG31676-GAL4* expression in PNs after intersecting with *GH146-Flp*. Similar numbers of PNs are labeled at 24h, 48h, and 72h APF. VA1d, but not DC3, is labeled, enabling us to map Cluster #3 to VA1d *(CG31676+)* and Cluster #4 to DC3 *(CG31676–)*. In addition, DA1 and DL3 are labeled. Therefore, the remaining *acj6−/CG31676+ cells* (Clusters #6 and #6’; see Figure S3 B) correspond to DL3 PNs. Scale bar, 20 µm. (G) Summary of the mapping of 6 PN classes (4 from the adPN lineage and 2 from the lPN lineage) to corresponding transcriptome clusters. Markers used for unambiguous mapping (Figures 3, 4A-D, and S3B, C) are listed. N-cadherin (Ncad) staining was used to label neuropil (red). See also Figure S3.

Using *Mz19-GAL4*, we mapped two *Mz19+/acj6*+ clusters to VA1d and DC3 in a two-to-two manner (Figure 3C; Clusters #3 and #4), but we were not able to resolve which cluster belonged to which PN class. VA1d and DC3 PNs are closely related: VA1d PNs are born immediately after DC3 PNs from the same anterodorsal neuroblast lineage (Yu et al., 2010), and the two classes target their dendrites to neighboring glomeruli, both postsynaptic to ORNs originating from trichoid sensilla (Couto et al., 2005). We sought to establish a one-to-one mapping between PN classes and transcriptome clusters by identifying markers that distinguish these two PN classes. We searched for differentially expressed genes and found a few that were robustly expressed in only one of the two clusters. Among them, *CG31676* exhibited the strongest differential expression (Figure 4D; compare Clusters #3 and #4).

To examine patterns of reporter expression driven by a *CG31676* driver, we generated *CG31676-GAL4* using a CRISPR/Cas9-based method (Diao et al., 2015). The transgene cassette, inserted into the first intron of *CG31676*, contained a splice acceptor (SA) sequence followed by T2A peptide sequence and the *GAL4* coding sequence (Figure 4E). This straightforward method generates a driver that is highly likely to recapitulate endogenous gene expression (Diao et al., 2015). After intersecting with *GH146-Flp, CG31676-GAL4* labeled a similar number of PNs at 24h, 48h, and 72h APF (Figure 4F). Notably, it labeled VA1d but not DC3 at both 48h and 72h APF (Figure 4F). This finding enabled us to unambiguously map the *Mz19*+/*acj6*+/*CG31676*+ Cluster #3 to VA1d PNs and the *Mz19+/acj6+/CG31676–* Cluster #4 to DC3 PNs.

In addition to DA1 (#2 and #2’) and VA1d (#3), *CG31676-GAL4* also strongly labeled DL3, which is targeted by *acj6−* lPNs (Figure 4F and S3A) (Jefferis et al., 2001). Among the 30 clusters, only two clusters (#6 and #6’) were *CG31676+/acj6−* (Figure 4D and S3B), and these two clusters display highly similar transcriptomes as reflected in their close proximity in the tSNE projection. We therefore mapped Clusters #6 and #6’ to DL3 PNs. *CG31676-GAL4* transiently labeled two other glomeruli, DL2a and DA4l (Figure S3A); both are targeted by *acj6*+ adPNs, but we could not unambiguously assign them to corresponding clusters.

Among the 6 glomerular classes we have mapped, four correspond to a single transcriptome cluster each, but DA1 and DL3 PNs each corresponded to two clusters (Figure 4D and Figure 4G). All PN classes are born in a stereotyped order within a specific lineage, and most PN classes are born consecutively within a single time window (Jefferis et al., 2001; Yu et al., 2010; Lin et al., 2012). Interestingly, however, DA1 and DL3 PNs are the only two exceptions: they are born in two time windows separated by more than 24 and 12 hours, respectively (Figure S3D; Lin et al., 2012). We speculate that this birth timing difference may contribute to the transcriptome heterogeneity of DA1 and DL3 PNs. For DA1 PNs, we found that *fruitless* (*fru*), encoding a transcription factor and a key regulator of male sexual behavior as a result of sex-specific alternative splicing (Baker et al., 2001; Dickson, 2008), was expressed only in the large cluster (#2). This is consistent with a previous finding that *NP21-GAL4* (GAL4 inserted into a *fru* intron near the sexually dimorphic splicing site) only labels DA1 PNs after intersecting with *GH146-Flp* (Kimura et al., 2005; Stockinger et al., 2005; Potter et al., 2010). On the other hand, *CG45263* was only expressed in the small cluster (#2’) (Figure S3C). For DL3 PNs that also matched with two clusters, we identified genes that were expressed only in one of the DL3 clusters, including a transcription factor *mirror (mirr)* and *CG7358* (Figure S3B). It will be interesting to investigate in the future whether the transcriptional differences between Clusters #2 and #2’ and between Clusters #6 and #6’ reflect only differences in birth timing, or potential differences in biological functions.

In summary, by using a combination of existing markers and new markers discovered using single-cell RNA-seq, we have unambiguously mapped 6 PN classes (those targeting to the DC2, DA1, VA1d, DC3, VM2, and DL3 glomeruli) to corresponding transcriptome clusters (Figure 4G). Our results demonstrate that the combination of genetic drivers and single-cell RNA-seq offers a simple strategy for mapping transcriptome clusters to cell types.

### Identification of New Lineage-specific Transcription Factors that Regulate Dendrite Targeting

Transcription factors (TFs) play important roles in determining cell fate and wiring specificity, including in *Drosophila* PNs (Komiyama et al., 2003; Zhu et al., 2006; Komiyama and Luo, 2007). Our single-cell transcriptome analysis identified TFs that were differentially expressed in different clusters. For example, Prospero was predicted to be expressed in a majority of PNs including all *Mz19*+ PNs, whereas Cut was predicted to be expressed in a few PNs, all of which were *Mz19–* (Figure S4A). Indeed, antibody staining validated these predictions (Figure S4B), and the expression of Cut is consistent with our previous finding (Komiyama and Luo, 2007).

In addition to *acj6* and *vvl*, single-cell transcriptome analysis also suggested several new TFs that were expressed in a lineage-specific manner. Specifically, *C15* and *knot* were predicted to be expressed only in adPNs and *unplugged (unpg)* was predicted to be expressed only in lPNs (Figure 5A). We confirmed these predictions by immunostaining using antibodies against C15 and Knot, and a *lacZ* reporter for *unpg* (Figure 5B). *knot* has been shown to play a critical role in controlling dendrite development of *Drosophila* class IV dendritic arborization (da) neurons (Jinushi-Nakao et al., 2007), and *unpg* is a marker for specific neuroblast sublineages in the *Drosophila* embryonic ventral nerve cord (Cui and Doe, 1995; Urbach and Technau, 2003). *C15*, encoding a homeobox-containing protein, is a homolog of human *Hox11* gene critical in regulating a gene network in the developing *Drosophila* leg (Campbell, 2005), but its role in the nervous system has not been explored. We tested whether *C15* plays a role in controlling PN dendrite targeting using loss-of-function and gain-of-function analyses.

**Figure 5.**
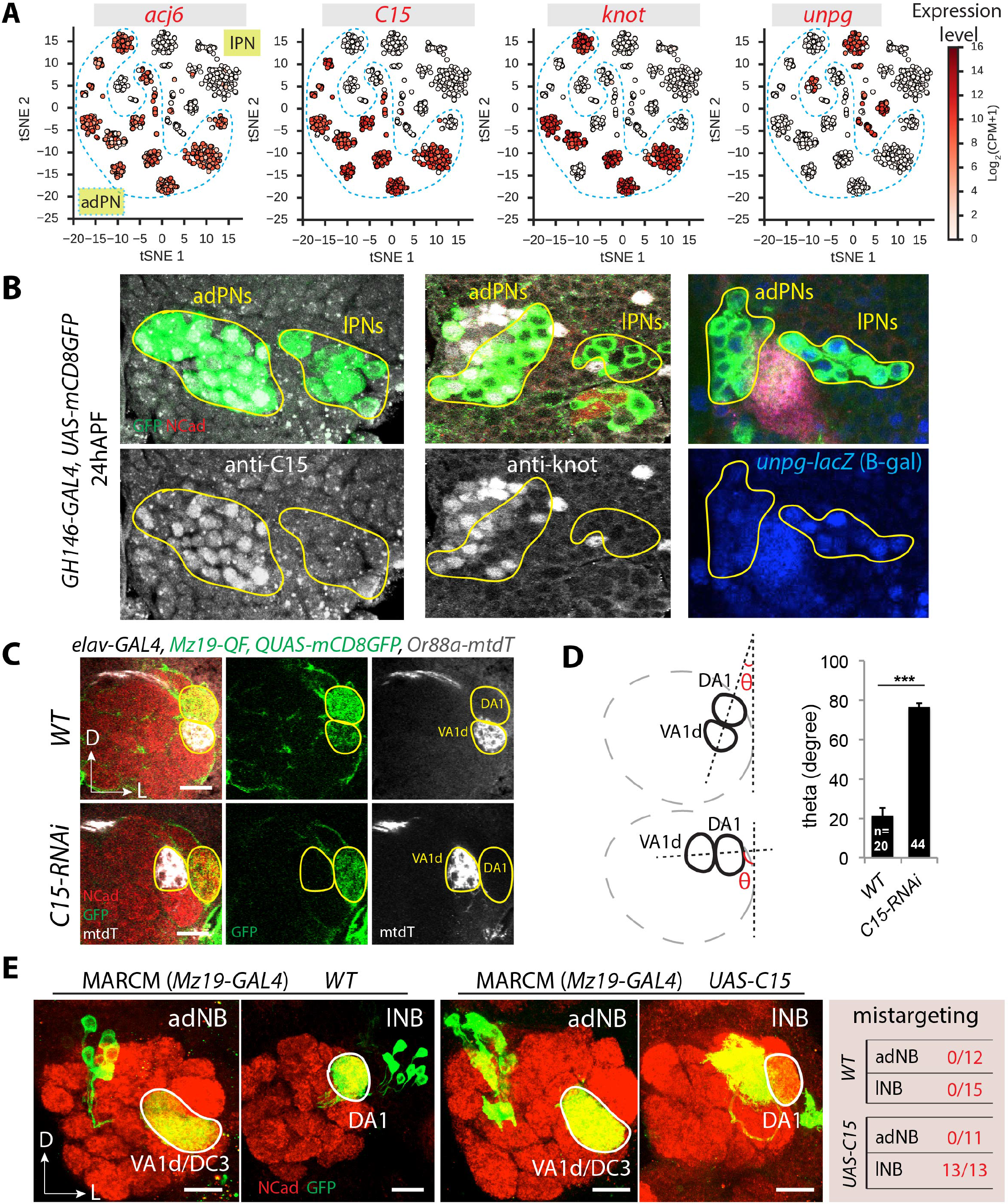
Identification of New Lineage-specific Transcription Factors from Single-cell RNA-seq. (A) Visualization of *GH146*+ PN cells using tSNE as in Figure 2E showing expression of *acj6, C15, knot*, and *unpg*. adPNs are outlined and remaining cells are lPNs (as determined based on *acj6* expression). *C15* and *knot* are only expressed in adPNs, and *unpg* is only expressed in lPNs. (B) Expression of C15 and Knot (antibody staining in white) in *GH146*+ PNs is restricted to adPNs *(GH146*+ cells are green, the adPN and lPN lineages are outlined in yellow). *unpg* (visualized using a lacZ reporter) is restricted to lPNs at 24h APF, consistent with RNA-seq data in (A). (C) Loss-of-function analysis of *C15* using *elav-GAL4* driven *UAS-C15-RNAi* (line #2, about 80% reduction of *C15* mRNA; see Figure S4C). *elav-GAL4* crossed with *w^1118^* flies were used as wild type (WT) control. When *C15* is knocked down, the VA1d glomerulus (visualized by VA1d ORN axons labeled by *Or88a-mtdT)* displays a dorsal shift. In addition, GFP signal in VA1d PN dendrites (visualized by *Mz19-QF* driven *QUAS-mCD8GFP)* is undetectable. Scale bar, 20 µm. (D) Quantification of position shift of the VA1d glomerulus due to C15 knockdown seen in (C). θ is the angle between a line drawn from the center of VA1d and DC3 and the vertical line demarcating the dorsoventral axis. Error bars are SEM. ***, P < 0.001 (t test). Sample numbers are indicated. (E) Gain-of-function analysis of *C15* in *Mz19-GAL4*+ overexpression MARCM clones. In *WT* control, dendrites of adPN neuroblast (adNB) clones target the VA1d and DC3 glomeruli and lPN neuroblast (lNB) clones target the DA1 glomerulus. When *C15* is misexpressed, dendrite targeting of adNB clones is not affected, whereas dendrite targeting of lNB clones is affected with 100% penetrance, as quantified at the right. Scale bar, 20 µm. N-cadherin (Ncad) staining was used to label neuropil (red). See also Figure S4.

For the loss-of-function experiment, we used a previously established RNAi-based genetic screening platform (Ward et al., 2015). Specifically, we used *elav-GAL4* to knockdown *C15* in all neurons, *Mz19-QF-driven QUAS-mCD8GFP* to monitor dendrite targeting of VA1d and DA1 PNs [note that *Mz19-QF* labels DA1 and VA1d, but not DC3 PNs in wild type (Hong et al., 2012)], and Or88a enhancer/promoter fused with myristolated tdTomato *(Or88a-mtdT)* to monitor axon targeting of VA1d ORNs. We first tested the efficiency of two *UAS-RNAi* lines against *C15* using qRT-PCR following pan-neural expression, and found that they reduced *C15* mRNA by 40% and 80%, respectively (Figure S4C). Pan-neuronal knockdown of *C15* using the stronger RNAi line caused a highly penetrant position shift of the VA1d glomerulus (visualized by *Or88a-mtdT*+ axons) to a dorsal- and medial-region without affecting DA1 dendrite targeting (Figure 5C, 5D, and S4D). We also observed the loss of dendrites in the VA1d glomerulus (visualized by *QUAS-mCD8GFP* driven by *Mz19-QF*). The loss of VA1d dendrites could be because (1) C15 controls the expression of *Mz19-QF* in VA1d PNs, (2) VA1d neurons are not born or die, or (3) VA1d dendrites mistarget to the DA1 glomerulus.

For the gain-of-function experiment, we used the *Mz19-GAL4*-based MARCM system to misexpress *C15*. In wild type, *Mz19*+ adPNs target to the VA1d and DC3 glomeruli and *Mz19*+ lPNs target to the DA1 glomerulus (Figure 5E, left panels; Figure 3B). However, when *C15* was misexpressed, *Mz19*+ lPNs (which are DA1 PNs only) sent their dendrites to regions outside the DA1 glomerulus, which included VA1d, DC3, DA3, and DA4l that are all normally targeted by adPNs, while *Mz19*+ adPNs still targeted dendrites correctly (Figure 5E, right panels; S4E and S4F). These data suggest that the transcription factor C15, as with Acj6 and Vvl (Komiyama et al., 2003), instructs lineage-specific PN dendrite targeting.

### Transcriptomes of Closely Related PN Classes Exhibit the Largest Differences during Circuit Assembly

How do neuronal transcriptomes change as development proceeds? By mapping clusters from single-cell RNA-seq data to specific PN classes at different developmental stages, we can address this key question at the resolution of single PN classes. Here, we focused our analyses on the three classes of *Mz19*+ adPNs and lPNs, which have been unequivocally mapped to specific transcriptome clusters (Figure 4D).

Following the coarse patterning of PN dendrites at 24h APF, ORN axons invade the antennal lobe beginning around 30h APF to identify their PN partners, until they match with cognate PNs and establish discrete glomerular compartments that are first visible around 48h APF (Jefferis et al., 2004). Following further expansion of terminal branches of ORN axons, PN dendrites, and synaptogenesis after 48h APF, pupae become adults at around 100h APF. Using the intersection of *Mz19-GAL4* and *GH146-Flp*, which labels three PN classes, we sequenced and analyzed 485 single cells from five time points (~100 cells each): 24–30h APF, 36–42h APF, 48-54h APF, 72–78h APF, and 1–2d adult (Figure 6A).

**Figure 6.**
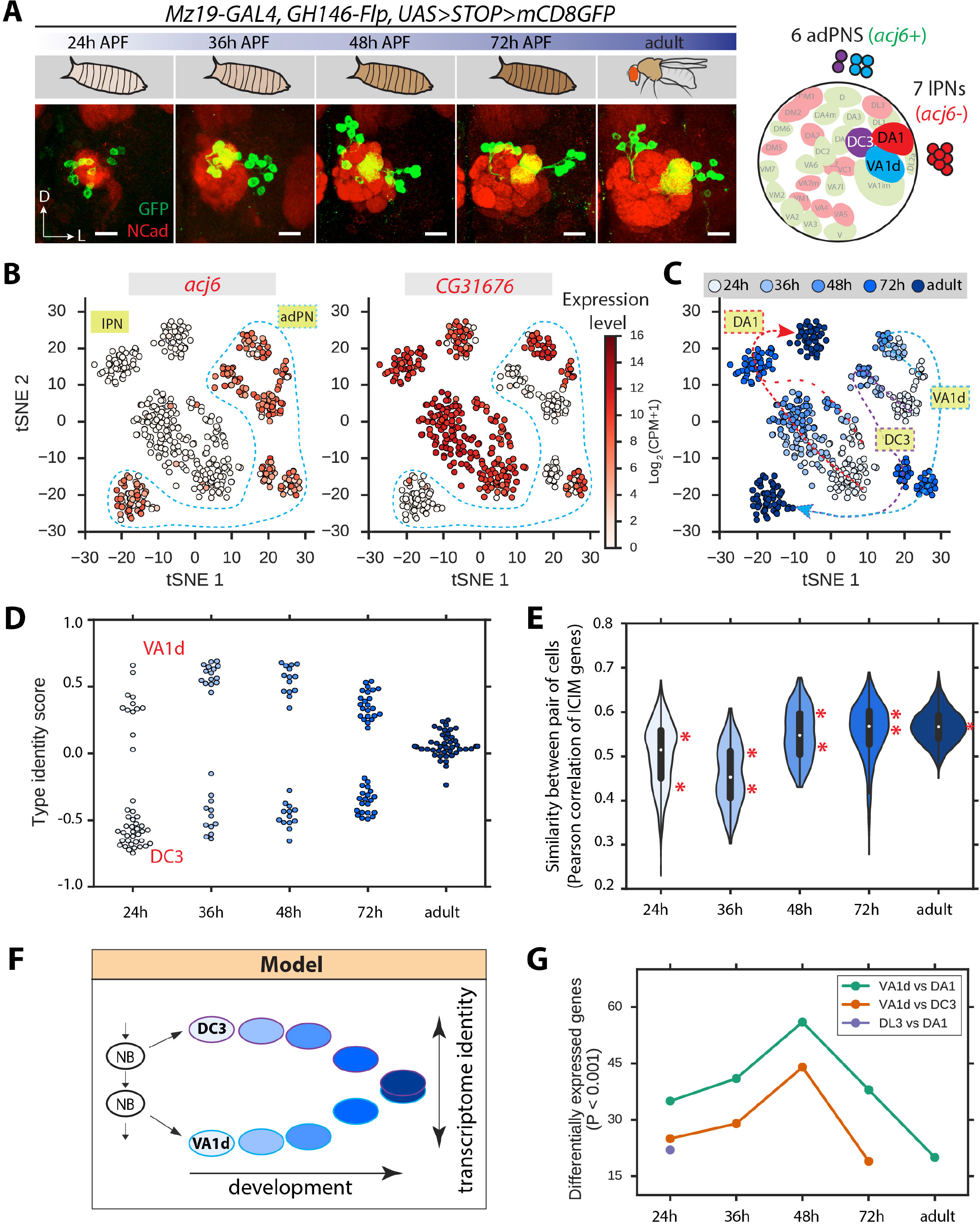
Transcriptome Analysis of *Mz19*+ PNs across Developmental Stages. (A) Representative confocal images (projections) of *Mz19*+ PNs *(Mz19-GAL4* intersected with *GH146-Flp)* from 5 different stages. Schematic (right panel) shows the three classes of *Mz19*+ PNs, which belong to the adPN (*acj6*+) and lPN lineages *(acj6−)*. N-cadherin (Ncad) staining was used to label neuropil (red). D, dorsal; L, lateral. Scale bar, 20 µm. (B) Visualization of *Mz19*+ PNs from developmental stages ranging from 24h APF to adult using tSNE based on 497 genes identified using ICIM. Color shows expression of *acj6* and *CG31676* (CPM, counts per million). *acj6*+ cells are adPNs (VA1d and DC3, outlined), and *acj6-* cells are lPNs (DA1). Within adPNs, *CG31676*+ cells are VA1d PNs, and CG31676− cells are DC3 PNs. Note that *CG31676* is turned off in all adult adPNs (see Figure 6C for developmental stages of these cells). (C) Visualization of *Mz19*+ PNs from different developmental stages, as in (B), with color indicating developmental stages. *acj6* and *CG31676* expression patterns (Figure 6B) enabled unambiguous identification of three PN classes, which are labeled. Dashed lines indicate the developmental trajectories of these classes. VA1d and DC3 PNs are distinct at all pupal stages, but merge to form one cluster in the adult. The densely and sparsely dashed red lines represent the trajectories of Cluster 2 and 2’ respectively, which become indistinguishable by 72h APF and remain so in the adult. (D) Type identity score of VA1d and DC3 PNs from five developmental stages. Each dot represents one cell. Colors show developmental stages, as in (C). The identity score is calculated as a scaled sum of the 23 most significantly differentially expressed genes between VA1d and DC3 PNs (P < 10^−5^; Experimental Procedures). 13 genes are highly expressed in VA1d PNs, but not in DC3 PNs, and these contribute positively to the score. 9 genes displayed the opposite expression pattern, and these genes contribute negatively to the score. Scores range from −1 to 1, where −1 indicates high expression of the DC3 signature genes and no expression of the VA1d signature genes, and +1 indicates the opposite expression profile. (E) Violin plot showing the distribution of transcriptome similarity between all pairs of *Mz19*+ adPNs. Transcriptome similarity was calculated as the Pearson correlation of expression levels of the 497 genes identified in an unbiased manner by using ICIM on *Mz19*+ PNs. Peaks are indicated by asterisks. The upper peak (higher similarity) consists of pairs of cells in which both cells belong to the same class (i.e., both VA1d or both DC3). The lower peak consists of pairs of cells in which the two cells belong to different classes (i.e., VA1d and DC3). In the adult, the distribution is unimodal, reflecting the dissipation of transcriptome differences between the two classes. (F) Schematic model of *Mz19*+ adPN development. VA1d and DC3 PNs are derived from a common neuroblast (NB) lineage. Our data indicate that the transcriptomes of these two PN classes are distinct at pupal stages and become indistinguishable in the adult. (G) Differentially expressed genes between PN classes belonging either to the same lineage (VA1d and DC3 from the adPN lineage, or DL3 and DA1 from the lPN lineage) or to different lineages (VA1d and DA1). Adult data does not exist for VA1d vs DC3 PNs, as they are indistinguishable at adult. For DL3 PNs, we only have data for 24h APF. See also Figure S5.

We performed clustering analysis using ICIM and tSNE, which revealed that *Mz19+/acj6*+ (VA1d and DC3) and *Mz19+/acj6−* (DA1) cells were clearly separable at all time points (Figure 6B). Interestingly, VA1d PNs and DC3 PNs formed distinct clusters at the four pupal stages, but merged into a single cluster in the adult (Figure 6B and 6C), suggesting that the transcriptomes of VA1d and DC3 PNs become more similar during cell maturation and ultimately converge on a common state in adulthood. To test this idea quantitatively, we calculated cell type identity scores using the 23 most differentially expressed (DE) genes (genes having differential expression P values < 10^−5^) between VA1d and DC3 PNs, and found that the difference between the transcriptional states of these two PN classes was maintained from 24h to 48h APF, but began to shrink at 72h APF, and in the adult the two classes were indistinguishable (Figure 6D). To quantify the similarity between the transcriptional states of VA1d and DC3 PNs in an alternative, unbiased genome-wide fashion, we calculated the Pearson correlation between the expression profiles of all pairs of cells based on 497 genes identified by ICIM. This analysis revealed a similar result, confirming that transcriptome differences between VA1d and DC3 PNs disappeared in the adult (Figure 6E). Indeed, clustering analyses using only adult VA1d and DC3 PNs failed to find distinct populations (data not shown). Collectively, these data indicate that VA1d and DC3 PNs, which are born consecutively from a common neuroblast in the adPN lineage and target dendrites to neighboring glomeruli, have peak transcriptome differences during early pupal stages (24–48h APF) when PNs are refining their dendrite targeting and presenting cues for partner ORNs to match. These differences progressively diminish in late pupal and adult stages, until the two PN classes are transcriptionally indistinguishable (Figure 6F).

These observations suggest that PN subtype identity genes, which distinguish VA1d and DC3 during the wiring stages, are down-regulated once wiring specificity is established. To test this idea, we systematically identified genes that were differentially expressed at different pupal and adult stages in all *Mz19*+ PNs. Gene ontology (GO) analysis indeed revealed that down-regulated genes consisted of factors associated with development and differentiation, whereas most up-regulated genes were associated with metabolic processes (Figure S5A). Clustering of genes based on their dynamic expression pattern revealed waves of gene expression consisting of genes that are coordinately turned down or up at different developmental stages (Figure S5B). Interestingly, many more TFs and cell-surface and secreted molecules (CSMs) were down-regulated than up-regulated (Figure S5B). Our RNA-seq data showed that *CG31676*, which is expressed in VA1d but not DC3 PNs at 24h APF (Figure 4D, G), was turned off in both PN classes in adult while its expression in DA1 persisted (Figure S5C); this was validated with *CG31676-GAL4* expression analysis across developmental stages (Figure S5D).

Next we asked whether transcriptomes of PN classes from the same neuroblast lineage are more similar than those from different lineages. We found that the transcriptome differences between VA1d and DC3 PNs (both adPNs) were consistently smaller than that between VA1d and DA1 PNs (adPNs and lPNs, respectively) across developmental stages (Figure 6G). Similarly, the transcriptome differences at 24h APF between DA1 and DL3 lPNs (we only have DL3 data at 24h APF) were similar to those between VA1d and DC3 but smaller than those between VA1d and DA1 PNs. All four PN classes target to adjacent glomeruli in the dorsolateral antennal lobe (Figure 6A, right). Thus, lineage origin appears to contribute more to transcriptome similarities than dendrite targeting position.

### PN Subtype Identity is Specified by a Combinatorial Molecular Code, Not Unique Subtype-specific Marker Genes

How is cell type identity encoded in the transcriptome? Cell type identity could be specified in two alternative ways: (1) each cell type expresses at least one unique gene, or (2) each cell type expresses a unique combination of genes. The strategy used for specifying cell type identity in the nervous system remains an unresolved question. To comprehensively address how neuronal subtype identity is encoded in PNs, we approximated the 30 *GH146*+ transcriptome clusters as 30 subtypes, and searched for marker genes that are uniquely expressed in a single subtype. We designed two criteria for identifying such marker genes: 1) the gene must be robustly expressed within a cluster (>7 CPM, or log_2_(CPM+1) > 3, in >50% of the cells of a cluster); 2) the gene must not be expressed in any other cluster (>7 CPM in <10% of the cells of any other cluster). Using these criteria, we identified only six unique marker genes (Figure S6A). These genes were sufficient to specify only 5 of the 30 *GH146*+ PN clusters. When we relaxed the search criteria (see Experimental Procedures), we quickly entered a regime where the identified genes were expressed in multiple clusters, indicating that they were not unique (Figure S6B).

We note that identification of unique marker genes is limited by technical artifacts such as transcript dropouts. We calculated the dropout rate as a function of expression level (Figure S6C), and based on this we estimate that the probability that we failed to detect marker genes in all 25 out of the 30 *GH146*+ PN subtypes due to dropout events is < 10^−62^ (Experimental Procedures). Moreover, the unique markers that we identified were found in clusters represented by few cells in our data set, rather than those having many cells, arguing further that dropout does not account for our inability to detect unique markers in most subtypes. Thus, with few exceptions, *GH146*+ PNs lack marker genes that uniquely specify subtypes.

Next, we sought to identify combinatorial molecular codes for cell type identity. We searched for a minimal set of genes that can uniquely specify PN subtypes using an information theory-based approach. We calculated the information content of each gene with respect to PN subtype identity, formally defined as the mutual information between the binarized expression state of the gene (ON/OFF) and PN cluster identity (Experimental Procedures). We ranked genes by their information content, and then selected a minimal set of genes by greedy search, iteratively drafting the gene carrying the most non-redundant information about identity into the set until 95% of the uncertainty of subtype identity was explained. The result of this search is a set of genes for which knowledge of their expression states (ON/OFF) alone is sufficient to classify subtype identity with high accuracy.

We first applied this strategy to the three *Mz19*+ PN classes that were unequivocally mapped to specific glomeruli. We found that only two genes, *C15* and *CG31676*, were sufficient to distinguish these three subtypes (Figure 7A). Based on these two genes alone, 92% of the uncertainty of classification of individual *Mz19*+ PN cells into subtypes was explained. Interestingly, both *C15* and *CG31676* were independently identified and characterized earlier in our study (Figure 4D and Figure 5). This finding demonstrates that this approach can identify sets of genes that robustly encode cell type identity in a combinatorial manner.

**Figure 7.**
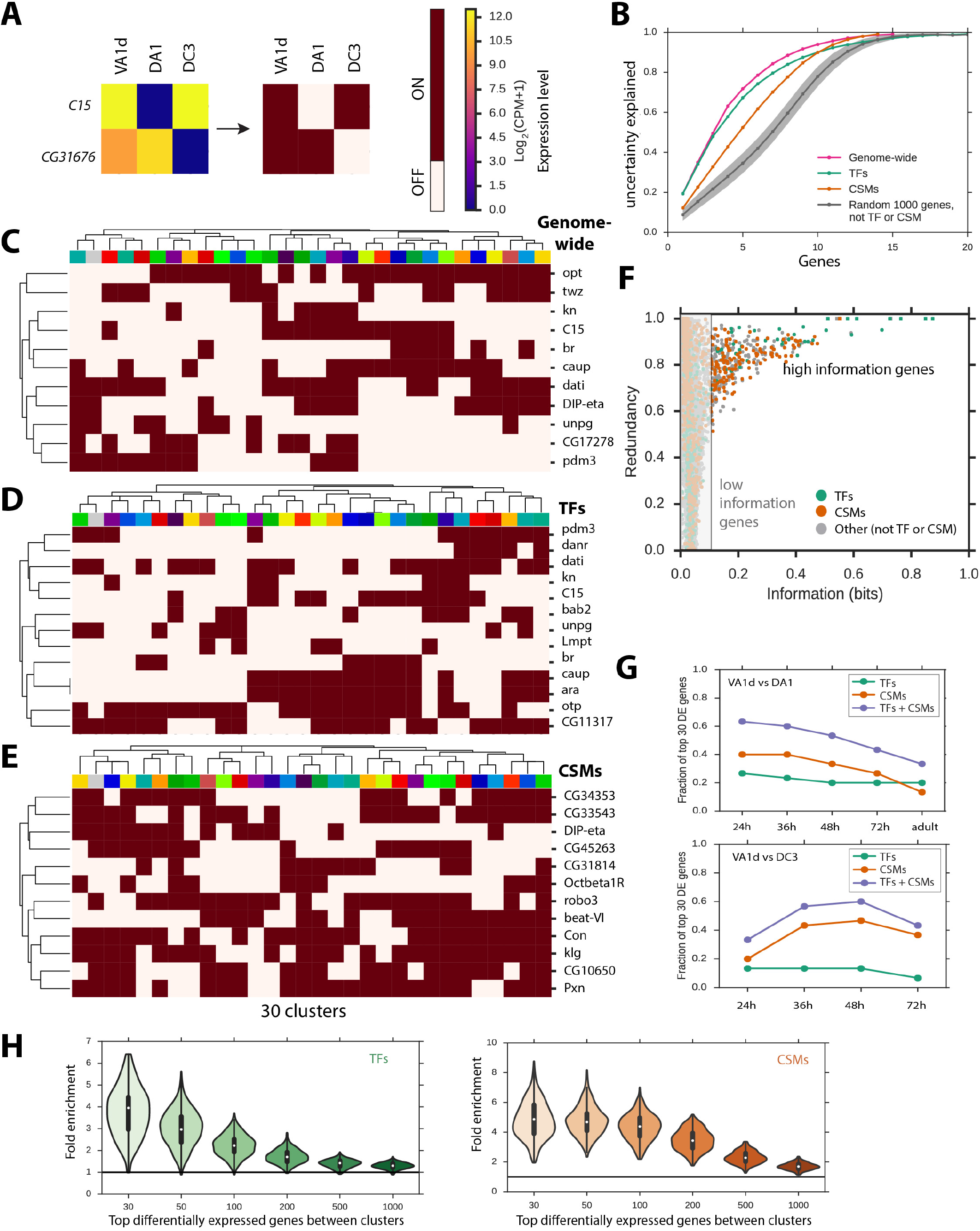
Combinatorial Molecular Codes of PN Subtype Identity. (A) Minimal combinatorial code for subtype identity among *Mz19*+ PNs identified using an information theory-based approach. Left panel shows median expression level of each gene among cells belonging to each *Mz19*+ class. Right panel shows binarized expression levels (ON/OFF) of the same genes (using a cutoff of 7 CPM, or log_2_(CPM+1) = 3). Each *Mz19*+ PN class expresses a distinct combination of these two genes. (B) Information contained in minimal combinatorial codes for *GH146*+ subtype identity of varying sizes and origins. Horizontal axis is the number of genes included in the code. Vertical axis is the amount of uncertainty (entropy) of cell type classification that is explained by the code. Colors denote codes constructed from different sets of genes. The genome-wide code (pink) is constructed from all genes, while the TF (green) or CSM (orange) codes use only 1048 TF or 1062 CSM genes. Gray denotes codes constructed from 1,000 randomly chosen non-TF and non-CSM genes, with the line indicating the median of and the shading indicating the standard deviation across 100 replicates, respectively. (C-E) Minimal combinatorial codes for *GH146*+ subtype identity constructed from (C) all genes, (D) TFs, or (E) CSMs. Heat map indicates the binary expression state of the gene in a cluster, where brown denotes the ON state and white denotes the OFF state (using a cutoff of 7 CPM, or log_2_(CPM+1) = 3). Clusters and genes are arranged by hierarchical clustering. (F) Redundancy and information content of individual genes with respect to *GH146*+ PN subtype identity. Each dot is a gene. Information of a gene is the mutual information of its binarized expression level and cluster identity. Redundancy of a gene is the fraction of its information that is contained in the minimal combinatorial code (Figure 7C). Genes are colored to indicate their type (TFs, green; CSMs, orange; other, gray). Squares indicate genes that are included in the minimal combinatorial code. (G) Representation of TFs and CSMs among the top 30 differentially expressed genes between *Mz19*+ PN subtypes. Top panel shows VA1d and DA1 cells, while bottom panel shows VA1d and DC3 cells. Vertical axis shows the fraction of the 30 most significantly differentially expressed genes that are TFs (green), CSMs (orange), or TF + CSM (blue) at each developmental stage. Adult stage is absent from the VA1d versus DC3 comparison because these two PN classes cannot be distinguished based on their transcriptomes. (H) Enrichment of TFs and CSMs among the top differentially expressed genes between pairs of clusters of *GH146*+ cells. Horizontal axis shows the number of significantly differentially expressed genes under consideration. Vertical axis shows the distribution of enrichment among pairs of 30 clusters (435 pairs). Enrichment is calculated relative to the genomic representation of TFs (6.8%) and CSMs (6.8%), which is indicated by the horizontal line. See also Figures S6 and S7.

We then applied this strategy to the 30 *GH146*+ PN subtypes described in this study. We identified 11 genes whose expression states uniquely identified every PN subtype (Figure 7C and Figure S7C). Knowledge of the expression states of these genes alone is sufficient to resolve 95% of the uncertainty in classification of individual *GH146*+ PN cells into subtypes (Figure 7B, pink line). While we used log_2_(CPM+1) = 3 as the threshold for binarization of expression state (ON/OFF), similar results were obtained using a range of different thresholds (Figure S7A, B). Together, these analyses demonstrate that *GH146*+ PN subtype identity can be specified using a combinatorial code composed of expression states of only 11 genes. This code is more compact than a code specifying each subtype using a unique marker (30 genes required), but substantially above the theoretical minimum of 5 genes (which can specify 2^5^ or 32 binary states).

### TFs and CSMs Redundantly Specify PN Subtypes and Are Highly Enriched Among Differentially Expressed Genes

What types of genes distinguish neuronal subtypes? It is widely thought that transcription factors (TFs) establish and maintain cell type identity, while cell-surface and secreted molecules (CSMs) determine wiring specificity. But there has not been, to our knowledge, genome-wide analysis to show this in an unbiased manner. Strikingly, 8 of the 11 genes in the minimal combinatorial code identified by our information theory-based analysis above were TFs (Figure 7C), supporting a central role for TFs in encoding cell type identity. To further explore the roles of TFs and CSMs in class identity, we searched for minimal codes for cell type identity consisting only of TFs or CSMs using our information theory-based approach. We used previously annotated lists of TFs (www.FlyTF.org) and CSMs (Kurusu et al., 2008), each containing about 1000 genes. We found minimal codes consisting of 13 TFs (Figure 7D and Figure S7C) or 12 CSMs (Figure 7E and Figure S7C). Each of these codes was sufficient to resolve 95% of the uncertainty in classification of *GH146*+ cells into PN subtypes (Figure 7B). That is, the *GH146*+ cells can be classified into PN subtypes based on the expression states of either 13 TFs alone or 12 CSMs alone. The compactness of these minimal codes was similar to that of the most compact code obtained in our genome-wide search, which had 11 genes (Figure 7C). Thus, PN subtypes can be redundantly encoded by TFs or CSMs. Similar results were obtained when we examined combinatorial codes based on gene expression levels after discretization into four states (Off, Low, Medium, High) instead of binary states (ON/OFF) (data not shown).

We next directly quantified the information content and redundancy of individual genes. To measure redundancy, we compared the information gained when the gene was added to the minimal combinatorial code consisting of 11 genes to the information carried by the gene individually when it is alone (its information content). Highly redundant genes added little information beyond that already carried by the code. We found widespread redundancy in the encoding of cell identity encompassing both TFs and CSMs. There were many highly informative genes that were highly redundant (Figure 7F). Among the 134 highly informative and redundant genes (which individually reduced uncertainty about cell type classification by >20%, but >50% of this information was already contained in the minimal code), TFs and CSMs were highly enriched (30 TFs and 54 CSMs, comprising 22% and 40% of these genes, respectively) (Figure 7F). This result indicates that many genes in the most compact combinatorial code could be substituted with other genes. Indeed, by including these genes alongside the compact code, one can design a robust code for which removing any particular gene alone does not impair the ability to classify cells, similar to the design of error-correcting codes for digital communication over noisy channels. The existence of redundancy among both TFs and CSMs suggests that error-correction mechanisms could be employed in specification of cell fate and wiring specificity.

To evaluate whether TFs and CSMs are particularly informative with respect to subtype identity, we measured the amount of information contained within minimal combinatorial codes built from other genes (not TFs or CSMs) chosen at random from the genome (sampling 1,000 genes at random with 100 replicates). Randomly chosen genes carried significantly less information than TFs or CSMs (Figure 7B). This was reflected in the lower amount of information carried by minimal codes of various sizes constructed from randomly chosen non-TF and non-CSM genes compared with codes constructed from TFs or CSMs. These findings indicate that, on average, TFs and CSMs carry more information about *GH146*+ subtype identity than other genes.

To test this idea further, we asked whether TFs and CSMs are enriched among genes that are differentially expressed among PN subtypes. We focused initially on *Mz19*+ adPNs and lPNs that have been unequivocally mapped to specific transcriptome clusters. Analyses of differential expression comparing VA1d versus DA1 PNs, and VA1d versus DC3 PNs, revealed that TFs and CSMs accounted for a large proportion of differentially expressed genes (Figure 7G). Representation of CSMs peaked during the circuit assembly state (24–48h APF), consistent with a role for differential expression of CSMs in determining wiring specificity. To comprehensively test whether this hypothesis holds for all *GH146*+ PNs, we analyzed differentially expressed genes separating every pair of 30 clusters, comprising 435 (30 x 29 / 2) pairs altogether. For every pair of clusters, TFs and CSMs were highly enriched among differentially expressed genes, with the strongest enrichment found among the most significantly differentially expressed genes (Figure 7H). This finding supports the notion that expression of TFs and CSMs plays key roles in establishing PN subtype identity and determining their wiring specificity.

## Discussion

In recent years, single-cell RNA-seq has emerged as a powerful technique to investigate cellular heterogeneity, discover new cell types, and identify cell type-specific markers (reviewed by Saliba et al., 2014; Johnson and Walsh, 2017). While widely used to profile mammalian including human cells, it has not been systematically applied to invertebrate genetic model organisms such as *Drosophila*. Here we establish a robust single-cell RNA-seq protocol for cells from *Drosophila* brains across pupal and adult stages. By focusing on olfactory projection neurons (PNs), one of the best characterized cell types in the nervous system, our analyses shed new light on the relationship between the transcriptome, neuronal cell type, and development.

Several lines of evidence support the sensitivity and reliability of our single-cell RNA-seq protocol. First, differential gene expression identified by single-cell RNA-seq is highly consistent with previous literature. For example, we found highly correlated expression of two male-specific RNAs (Meller et al., 1997) at the level of individual cells (Figure S2A), and mutually exclusive expression of two lineage-specific transcription factors (Figure S2F) as previously reported (Komiyama et al., 2003). Second, we experimentally validated five differentially expressed transcription factors predicted from single-cell RNA-seq data (Figure 5A, B; Figure S4A, B). Third, sequencing of cells marked by known or newly identified PN class-specific markers matched well with specific transcriptome clusters (Figure 3; Figure 4), which not only provided validation of the transcriptome analysis, but also helped us to match transcriptome clusters with glomerular classes. Indeed, the protocol is sensitive enough to capture transcriptome differences within the same glomerular classes of PNs separated by birth timing (Figure S3). The fact that we have validated differential expression of many transcription factors, considered relatively less abundantly expressed (Ghaemmaghami et al., 2003), further support the sensitivity of our protocol. We note that the reliability of generating full-length cDNA material from single glia (30%) was not as high as from single neurons (90%; Figure S1D), suggesting that individual glia may contain fewer mRNA molecules, or the dissociation and sorting procedure may cause more damage to glia than to neurons. We expect that this approach can be generally applied to single-cell transcriptome analysis of many tissues and developmental stages in *Drosophila* and other organisms with small cell size, thus expanding the use of single-cell transcriptomics for addressing diverse biological questions.

We have developed a machine-learning algorithm called ICIM for unbiased identification of genes that distinguish subtypes. Because this algorithm recursively examines finer-grained subpopulations, it is capable of detecting genes that distinguish small subpopulations. This approach is conceptually similar to previously described iterative analysis methods (Usoskin et al. 2014; Zeisel et al. 2015; Tasic et al. 2016; Gokce et al. 2016). However, our approach may discriminate highly similar cell types with greater sensitivity than methods based on PCA (Usoskin et al. 2014; Tasic et al. 2016; Gokce et al. 2016) because it reduces the feature space to only those genes that are informative for distinguishing cell types. Use of this approach allowed us to distinguish 30 clusters of *GH146*+ PNs. Our classification is limited by sampling depth. While *GH146-GAL4* labels 40 PN classes, at least 17 of these classes contain only 1 cell per hemisphere. Many of our 30 clusters likely correspond to a single glomerular class. However, some of these clusters must contain cells representing more than one class. Sequencing of many more cells may resolve these classes into distinct clusters, resulting in a more complete description of PN transcriptome diversity.

Our analyses of transcriptome changes of identified PN classes across developmental stages provide, to our knowledge, the first demonstration that transcriptomes of neuronal subtypes exhibit the largest difference during development, coincident with circuit assembly. This could be because two key determinants for different PN classes are 1) from which neurons they receive information, and 2) to which neurons they send information. Once different PN classes establish differential connectivity during development, they may use largely the same neuronal machineries—such as neurotransmitter receptors to receive input from cholinergic ORNs and GABAergic or glutamatertic local interneurons as well as neurotransmission apparatus to send output (Wilson, 2013)—to convey different olfactory information in adults. Indeed, in the case of VA1d and DC3 PNs, even though they have clearly distinct transcriptomes during development, their transcriptomes become indistinguishable in adults (Figure 6). This finding has important implications for using single-cell RNA-seq to classify neuronal types, since most studies have exclusively focused on adults (e.g., Johnson et al., 2015; Usoskin et al., 2015; Zeisel et al., 2015; Foldy et al., 2016; Fuzik et al., 2016; Shekhar et al., 2016; Tasic et al., 2016); however, see (Darmanis et al., 2015; Hanchate et al., 2015; Pollen et al., 2015)]. While these studies have been highly successful in classifying major neuronal types, some functionally distinct subtypes may have been overlooked, resulting in an under-estimate of the diversity of neuronal types.

Transcription factors (TFs) and cell-surface and secreted molecules (CSMs) are widely considered to be key determinants of cell fate and wiring specificity, respectively. Our single-cell transcriptome analyses provided several objective criteria to support these notions. First, TFs and CSMs together account for more than 50% of the top 30 differentially expressed genes, whether between identified PN classes (Figure 7G) or among all GH146+ clusters (Figure 7H). Second, information theory-based analyses revealed that among the top 11 information-rich genes for distinguishing different PN subtypes, 8 are TFs. Third, TFs or CSMs alone contain nearly as much information in distinguishing different PN subtypes as all genes (Figure 7B). The TF and CSM codes are redundant in distinguishing different PN subtypes, consistent with the notion that differential expression of CSMs is a consequence of differential expression of TFs. Indeed, a key readout of TFs in determining neuronal subtype may be the control of differential expression of CSMs, such that different subtypes differentially respond to a common extracellular environment to achieve their wiring specificity. Supporting a role for TFs in regulating wiring specificity, we show that a newly identified lineage-specific TF, the homeobox-containing C15, can instruct lineage-specific dendrite targeting (Figure 5).

Finally, our analyses of PN transcriptomes shed light on the nature of the coding strategies that distinguish closely related neuronal subtypes. We found a very small number of markers that are uniquely expressed in single PN subtypes; instead, the identity of PN subtypes is largely determined by a combinatorial code that utilizes a number of genes between the number of subtypes and the theoretical minimum for a maximally compact code, suggesting redundancy. Our analyses reveal that highly informative genes in distinguishing PN subtypes are also highly redundant, such that the loss of any individual gene alone can in theory be compensated by remaining genes with little impact on PN subtype identity. We found that the transcriptomes of closely related PN classes, which become indistinguishable in adults, differed substantially during development (Figure 7G), consistent with a recent report showing that closely related retinal cells have dozens of differentially expressed CSMs (Tan et al., 2015). These findings may potentially explain why almost all previously identified genes that regulate PN dendrite targeting yielded only partially penetrant loss-of-function phenotypes even when null alleles were used. Such redundancy can provide robustness to wiring precision, and at the same time creates challenges for dissecting genetic control of wiring specificity using single gene manipulation (Hong and Luo, 2014). The transcriptomes of identified PN classes can inform design of more precise experiments in which simultaneous manipulation of multiple genes through loss- and gain-of-function approaches allows experimental testing of the combinatorial TF and CSM codes.

## Experimental Procedures

### Fly Stocks

The following fly lines were used in this study: *elav-GAL4* (Bloomington *Drosophila* stock center, BL#458), *Mz19-GAL4* (BL#34497) (Jefferis et al., 2004), *C15-RNAi* (line1, BL#27649), *C15-RNAi* (line2, BL#35018), *Alrm-GAL4* (BL# 67032). *Mz19-QF* (Hong et al., 2012), *GH146-GAL4* (Stocker et al., 1997), *UAS-STOP-mCD8GFP* (Hong et al., 2009), *GH146-Flp* (Potter et al., 2010), *unpg-lacZ* (Doe, 1992), *trol-GAL4* (NP5103-GAL4, Kyoto Stock Center #113584), *UAS-C15* (gift from Dr. Gerard Campbell) (Campbell, 2005), and *91G04-GAL4* (gift from Gerry Rubin) (Jenett et al., 2012).

*CG31676-GAL4* was generated using CRISPR/Cas9 based insertion of *SA-T2A-GAL4* into the first intron of CG31676 gene following the method described by Diao et al. (2015). In brief, a 2.5kb DNA fragment, containing a PAM site in the middle, within the first intron of the *CG31676* gene was PCR amplified from wild-type genomic DNA, and inserted into Blunt TOPO vector (Invitrogen). Then, *SA-T2A-GAL4* was PCR amplified from the *pT-GEM(1)* plasmid (addgene# 62893) and was assembled (NEBuilder HiFi DNA assemble kit) into TOPO-CG31676-intron construct three-nucleotide before the PAM site of the intron. This construct and gRNA plasmid *(pU6-BbsI-ChiRNA*, addgene# 45946) containing 20-nt target sequence before the PAM inserted into the BbsI site were co-injected to *nos-Cas9* (gift from Dr. Ben White) (Diao et al., 2015) embryos to obtain transgenic flies.

### MARCM Analysis

*hsFlp* based MARCM analyses were performed as previously described (Lee and Luo, 1999; Jefferis et al., 2001; Komiyama et al., 2004). *Mz19-GAL4* was used to label VA1d, DC3 and DA1 PNs. Larvae (24h to 48h after hatching) were heat shocked for 1h at a 37^o^C water bath. Both single cell and neuroblast clones could be observed in this fashion.

### Immunostaining

Tissue dissection and immunostaining were performed following previously described methods (Wu and Luo, 2006). Primary antibodies used in this study include rat anti-DNcad (DN-Ex #8; 1:40; DSHB), mouse anti-Prospero (1:200; DSHB), mouse anti-Cut (2B10; 1:50; DSHB), mouse anti-β-gal (1:500; Promega), chicken anti-GFP (1:1000; Aves Labs), rabbit anti-DsRed (1:250; Clontech), mouse anti-ratCD2 (OX-34; 1:200; AbD Serotec), rat anti-C15 (1:200; gift from Dr. Gerard Campbell) (Campbell, 2005), and guinea pig anti-knot (1:200; gift from Dr. Adrian Moore) (Jinushi-Nakao et al., 2007). Secondary antibodies were raised in goat or donkey against rabbit, mouse, rat, and chicken antisera (Jackson Immunoresearch), conjugated to Alexa 405, 488, FITC, Cy3, Cy5, or Alexa 647.

### Quantitative PCR (qPCR)

Total RNA from 3–5 day old adult fly heads was extracted using MiniPrep kit (Zymo Research, R1054). Complementary DNA was synthesized using an oligo-dT primer. qPCR was performed on a Bio-Rad CFX96 detection system. Relative expression was normalized to *Actin5C*.

*Actin5C* (F): 5’-CTCGCCACTTGCGTTTACAGT-3’

*Actin5C* (R): 5’-TCCATATCGTCCCAGTTGGTC-3’

*C15* (F): 5’-AGCGCTTCCACAAGCAAAAG-3’

*C15* (R): 5’-CCGTCTGTCGTCTCCACTTG-3’

### Imaging and Quantification Procedure

Confocal images were collected with a Zeiss LSM 780 and processed with ImageJ and Adobe Illustrator. For quantification of the angles in Figure 5C, the vertical line was drawn based on the position of two antennal lobes and the intersecting line was drawn through the centers of gravity of the VA1d and DA1 glomeruli, then the intervening angle was measured using ImageJ.

### Single-Cell RNA-sequencing

*Drosophila* brains with mCD8GFP-labeled cells using specific GAL4 drivers were manually dissected, and two optical lobes were removed. Single-cell suspension was prepared following Tan et al. (2015) with several modifications. Detailed protocol is available upon request. Single, labeled cells were sorted using Fluorescence Activated Cell Sorting (FACS) into individual wells of 96-well plates containing lysis buffer. Full-length poly(A)-tailed RNA was reverse transcribed and amplified using PCR following the SMART-seq2 protocol (Picelli et al., 2014) with several modifications. To increase cDNA yield and detection efficiency, we increased the number of PCR cycles to 25. To reduce the amount of primer dimer PCR artifacts, we digested the reverse transcribed first-strand cDNA using lambda exonuclease (New England Biolabs) (37^o^C for 30 min) prior to PCR amplification. Sequencing libraries were prepared from amplified cDNA using tagmentation (Nextera XT). Sequencing was performed using the Illumina Nextseq platform with paired end 75 bp reads.

### RNA-seq Data Analysis

We first provide an overview of our methods, then describe how these methods were applied to create each figure. All analysis was performed in Python using Numpy, Scipy, Pandas, scikit-learn, and a custom single-cell RNA-seq module. Sequencing reads will be deposited in the Sequence Read Archive. Preprocessed sequence data and code for analysis will be deposited to Data Dryad.

**Sequence alignment and preprocessing.** Reads were aligned to the *Drosophila melanogaster* genome (r6.10) using STAR (2.4.2a) (Dobin et al., 2013) with the ENCODE standard options, except “–outFilterScoreMinOverLread 0.4 –outFilterMatchNminOverLread 0.4 –outFilterMismatchNmax 999 –outFilterMismatchNoverLmax 0.04”. Uniquely mapped reads that overlap with genes were counted using HTSeq-count (0.7.1) (Anders et al., 2015) with default settings except “-m intersection-strict”. Cells having fewer than 300,000 uniquely mapped reads were removed. To normalize for differences in sequencing depth across individual cells, we rescaled gene counts to counts per million (CPM). All analyses were performed after converting gene counts to logarithmic space via the transformation Log_2_(CPM+1). Cells that were labeled with neuron-specific GAL4 drivers *(C155+, GH146+, Mz19+, 91G04+, Trol*+, and *CG31676*+ cells) were filtered for expression of canonical neuronal genes (*elav, brp, Syt1, nSyb, CadN*, and *mCD8-GFP*), retaining only those cells that expressed at least 4/6 genes at >15 CPM.

**Dimensionality reduction and clustering analyses using PCA/tSNE**. Single-cell RNA-seq yields high dimensional gene expression data. To visualize and interpret these data, we obtained two-dimensional projections of the cell population by first reducing the dimensionality of the gene expression matrix using principal component analysis (PCA), then further reducing the dimensionality of these components using t-distributed Stochastic Neighbor Embedding (tSNE) (van der Maaten and Hinton, 2008). We note that tSNE is a nonlinear embedding that does not preserve distances, so one cannot interpret the distances in the projected space as distances between gene expression profiles (i.e., the pre-transformation space).

We performed PCA on a reduced gene expression matrix composed of the top 500 overdispersed genes (as described below). To identify significant principal components (PCs), we examined the distribution of eigenvalues obtained by performing PCA after shuffling the gene expression matrix (with 100 replicates). A PC was considered significant if the magnitude of its associated eigenvalue exceeded the maximum magnitude of eigenvalues observed in the shuffled data. Significant components (typically 7–12 PCs) were used for further analysis. We further reduced these components using tSNE to project them into a two-dimensional space.

**Dimensionality reduction and clustering analyses using ICIM**. ICIM is an unsupervised machine learning algorithm that identifies a set of genes which distinguishes transcriptome clusters, which may correspond to cell types (described below). In our analysis of *GH146*+ PNs, this set typically includes ~500 genes. To visualize and interpret the single-cell gene expression data, we further reduced its dimensionality using tSNE to project the reduced gene expression matrix (consisting of only the genes identified by ICIM) into a two-dimensional space.

**Overdispersion analysis**. Genes that are highly variable within a population often carry important information for distinguishing cell types. We were interested in identifying such genes and using them for dimensionality reduction and clustering analyses. Variability of gene expression depends strongly on the mean expression level of a gene. This motivates the use of a metric called dispersion, which measures the variability of a gene’s expression level in comparison with other genes that are expressed at a similar level. Overdispersed genes are those that display higher variability than expected based on their mean expression level.

To identify overdispersed genes, we binned genes into 20 bins based on their mean expression across all cells. We then calculated a log-transformed Fano factor D(x) of each gene x D(x)

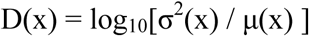

where σ^2^(x) is the variance and µ(x) is the mean of the expression level of the gene across cells. Finally, we calculated the dispersion d(x) as the Z-score of the Fano factor within its bin

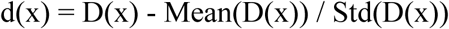

where Mean(D(x)) is the mean log-transformed Fano factor within the bin and Std(D(x)) is the standard deviation of the log-transformed Fano factor within the bin. We then rank genes by their dispersion and select the top genes for downstream analysis.

**Iterative Clustering for Identifying Markers (ICIM)**. To identify subpopulations of cells corresponding to PN subtypes, we developed an unsupervised machine-learning algorithm, which we call ICIM. We observed that standard dimensionality reduction and clustering methods using PCA and tSNE failed to discriminate subpopulations that corresponded to known PN lineages and molecular features. We attributed the failure of these methods to the high degree of similarity of transcriptional states among PN subtypes, which represent closely-related neurons having similar functions. All PN subtypes are born from one of the two common progenitor cells (neuroblasts) and have similar functional roles in the adult fly. Thus, PN subtypes may be distinguished by a small number of genes.

In the language of machine learning, the performance of dimensionality reduction and clustering methods depends critically on feature selection. Selection of informative genes that vary among cell types can improve discrimination in dimensionality reduction and clustering analysis.

We developed ICIM as a strategy to identify the most informative genes for distinguishing subpopulations within a population of closely-related cells in an unbiased way. Starting with a population of cells, we first identify the top 100 overdispersed genes within this population. Next we expand this set of genes by finding genes whose expression profiles are strongly correlated with the overdispersed genes (Pearson correlation > 0.5). We also filter this set of genes by (1) removing those having fewer than 2 correlated partners, and (2) those that are expressed in >80% of cells. Filter (1) removes noisy genes based on the idea that genes that carry information about cell type are expressed within gene modules and therefore have expression profiles that are correlated with at least one other gene. Filter (2) removes housekeeping genes that are detected in nearly all cells, and have variation in expression levels due to biological and technical noise, but this variation is not informative for purposes of distinguishing cell types. Cells are then clustered based on their expression profiles of these genes (average-linkage clustering using correlation metric). We cut the dendrogram at the deepest branch and partition the population into two subpopulations. The same steps are then performed iteratively on each subpopulation. Iteration continues until a population cannot be split into subpopulations because it is “homogeneous”. The termination condition is defined as the minimum terminal branch length (the most similar nearest-neighbor correlation distance between the expression profiles of cells) being larger than 0.2. This condition arises when the algorithm attempts to discover genes within a homogeneous population and finds a very large number of genes (typically >1000 genes) that vary in an incoherent manner between cells. When the algorithm terminates, we collect all genes that were identified at any stage. The result of this analysis is a set of genes that discriminate subpopulations within a population, which can be used for dimensionality reduction (as described above). We note that this algorithm identifies informative genes in an unbiased manner without knowledge of the ground truth of the number of cell types and their differences. The results of the algorithm were robust across a wide range of parameters.

**Differential expression analyses**. To find differentially expressed genes, we used the Mann-Whitney U test, a non-parametric test that detects differences in the level of gene expression between two populations. The Mann-Whitney U test is advantageous for this application because it makes very general assumptions: (1) observations from both groups are independent and (2) the gene expression levels are ordinal (i.e., can be ranked). Thus the test applies to distributions of gene expression levels across cells, which rarely follow a normal distribution. Using the Mann-Whitney U test, we compared the distributions of expression levels of every gene separately. P values were adjusted using the Bonferroni correction for multiple testing. Different significance thresholds for determining whether a gene is differentially expressed were used for various analyses in this work.

**Single-cell analyses to distinguish neurons and glia in Figure 1.** We formed a population consisting of 946 *GH146-GAL4*+ cells, 12 *elav-GAL4*+ cells, and 67 *alrm-GAL4*+ cells. We performed dimensionality reduction and clustering analysis using PCA and tSNE as described above. We identified the top 500 overdispersed genes in the population. We used PCA to reduce dimensionality, retaining 10 significant PCs. Then we projected the population into a twodimensional space using tSNE with perplexity 30 and learning rate 500 (Figure 1E). We also performed hierarchical clustering using complete linkage and a Euclidean metric based on manually selected neuronal and glial marker genes (Figure 1D).

**Removal *GH146*+ vPNs and APL neurons.** We initially formed a population consisting of 946 *GH146*+ cells. Using ICIM, we identified 158 genes that distinguish subtypes. We then projected the population into a two-dimensional space using tSNE. We observed several distinct subpopulations corresponding to *GH146*+ neuronal types that do not belong to the adPN or lPN lineages. Specifically, two clusters were composed of ventral PNs (vPNs), which robustly express several specific markers (*Gad1, Lim1*, and *toy*). Three other clusters were composed of APL neurons, which robustly express other specific markers (*Wnt4, VGlut*, and *fd102C*), and are arranged adjacent to one another by tSNE, reflecting the similarity of expression profiles among these cells. For subsequent analyses of *GH146*+ adPNs and lPNs, we removed these cells by excluding cells expressing 2/3 of these marker genes at >15 CPM.

**Single-cell analyses to distinguish PN subtypes among *GH146*+ cells in Figure 2.** We initially attempted to identify distinct subpopulations representing PN subtypes using PCA and tSNE for the 946 *GH146*+ PNs (including vPNs and APL neurons). We began by identifying the top 500 overdispersed genes and performing PCA to reduce the gene expression data to 10 significant PCs. Then we projected the population into a two-dimensional space using tSNE with perplexity 30 and learning rate 500. We observed that this analysis fails to separate distinct subpopulations (Figure 2B).

We next attempted to distinguish subpopulations corresponding to PN subtypes using ICIM and tSNE. Using ICIM (Figure 2C), we identified 561 genes for the 902 *GH146*+ PNs (after removing vPNs and APL neurons as described above). We projected these cells into a twodimensional space using tSNE using as a distance matrix the pairwise Pearson correlation of the expression profiles of these genes and perplexity 10, learning rate 250, and early exaggeration 4.0 (Figure 2D). We classified cells into clusters in an unbiased manner using HDBSCAN with min_cluster_size=5 and min_samples=3. Because tSNE computes a nonlinear embedding that does not preserve distances in the original space, the distances between cells cannot be directly interpreted in terms of similarity of expression profiles. As a consequence, there are cases where cells belonging to the same cluster are separated by larger distances than cells belonging to different clusters.

**Matching clusters to PN classes using known markers in Figure 3.** We formed a population consisting of 902 *GH146*+ cells, 123 *Mz19*+ cells, and 23 *91G04*+ cells. Using the 561 genes identified using ICIM on *GH146*+ cells, we projected this population into a two-dimensional space using tSNE with perplexity 15 and learning rate 1000. For visualization, we colored the cells according to their genotype, revealing that the *Mz19*+ and *91G04*+ cells belong exclusively to 5 classes (Figure 3C).

**Matching clusters to PN classes using newly identified markers in Figure 4.** We formed a population consisting of 902 *GH146*+ cells and 41 *trol-GAL4*+ cells. *trol-GAL4*+ cells that were not expressing *trol* (CPM < 7) were removed. Using the 561 genes identified using ICIM on *GH146*+ cells, we projected this population into a two-dimensional space using tSNE with perplexity 15 and learning rate 1000. For visualization, we colored the cells according to their genotype, revealing that the *trol*+ cells belong exclusively to 1 class (Figure 4C).

**Analyses of transcriptome changes during development in Figure 6.** To understand how transcriptional state changes during development and maturation of PN subtypes, we collected *Mz19*+ cells from flies at 5 stages of development: 24h, 36h, 48h, and 72h after puparium formation (APF), and 1–2 day adults. We formed a population consisting of 123, 83, 92, 92, and 95 cells after filtering to remove low quality cells and those not expressing neuronal markers. Using ICIM, we identified 497 genes that distinguish cell types and developmental stages. We projected this population into a two-dimensional space based on these genes using tSNE with perplexity 10 and learning rate 500. Cells formed several distinct subpopulations corresponding to different PN subtypes and developmental stages (Figures 6B and 6C). We assigned subpopulations to subtypes (DA1, VA1d, and DC3) based on the expression of key lineage factors (Figure 6B).

To quantify transcriptome changes in the closely related PN subtypes VA1d and DC3 during development, we devised a metric called the type identity score, which is the scaled sum of expression levels of genes that distinguish VA1d and DC3 cells. We identified these genes using differential expression analysis comparing VA1d and DC3 populations at all times that the two populations are distinct as determined by ICIM and tSNE (24h, 36h, 48h, and 72h APF). 78 cells were included in the VA1d group and 64 cells were included in the DC3 group. This analysis yielded 23 genes of which 13 are highly expressed in VA1d and 9 are highly expressed in DC3 cells at a significance level of P < 10^−5^ after the Bonferroni adjustment for multiple testing. We rescaled expression levels of these genes to the range 0 to 1 (by dividing each expression level by the maximum among the population), then calculated the type identity score I of each cell as the mean normalized expression level,

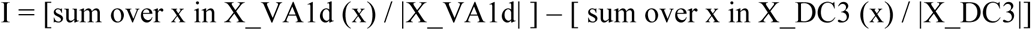

where X_VA1d is the set of genes that are highly expressed in VA1d and X_DC3 is the set of genes that are highly expressed in DC3, and |X_VA1d| and |X_DC3| are the cardinalities of these sets respectively. We then plotted the type identity scores of each cell at each developmental stage (Figure 6D).

As an alternative method to analyze transcriptome differences, we also examined correlations in transcriptome states in an unbiased genome-wide manner. This method has the advantage that it does not require the choice of a P value cutoff for determining significance. We formed a population consisting of *Mz19*+ adPNs (belonging to both subtypes VA1d and DC3) at each stage of development. Then we calculated the Pearson correlation of the expression profiles of the 497 genes identified by ICIM for every pair of cells (Figure 6E). These plots revealed a bimodal distribution consisting of two distinct peaks, which correspond to pairs of cells that both belong to the same subtype (more similar) and pairs of cells which belong to different subtypes (more distinct). As development proceeds, the transcriptome similarity of these two subtypes diminishes until vanishing, as reflected in the merging of these two distinct peaks and the emergence of a unimodal distribution in adulthood.

Finally, to characterize transcriptome changes distinguishing PNs in the wiring stages of development from PNs in adulthood, we performed differential expression analysis comparing the population of *Mz19*+ PNs at 24h APF to the population of *Mz19*+ PNs in adults. We found 1097 differentially expressed genes at significance level of P < 10^−5^ after the Bonferroni adjustment for multiple testing. This included 592 genes that were highly expressed in 24h APF cells, and 478 genes that were highly expressed in adult cells. We performed Gene Ontology (GO) analysis on these genes using Flymine and removed the redundant GO terms using REVIGO (Supek et al., 2011). We report the number of genes corresponding to and the P value of enrichment of each term (Figure S5A).

**Characterizing genes that distinguish PN subtypes in Figure 7.** To identify genes that distinguish closely related PN subtypes, we performed differential expression analysis comparing the *Mz19*+ VA1d and DA1 populations and the *Mz19*+ VA1d and DC3 populations at each developmental stage. Because the significance level of expression level differences depends on the number of cells involved in the comparison and the number of cells varies across developmental stages, we analyzed the top 30 differentially expressed genes regardless of their significance level. We note that the significance values of these genes were nearly all P < 10^−4^. We calculated the fraction of these genes that are transcription factors (TFs) or cell surface molecules (CSMs) based on the lists given in (www.FlyTF.org) or (Kurusu et al., 2008) (1048 TFs, 1062 CSMs) (Figure 7G).

We next performed a similar differential expression analysis comparing all pairs of subtypes. For each subtype, we formed a population consisting of *GH146*+ cells belonging to that subtype at 24h APF. For each pair of subtypes we calculated differential expression for each gene and ranked the genes by their significance level (P value). We then calculated the fraction of TFs or CSMs among the top N genes for varying N from 30 to 1000. We calculated the enrichment of TFs or CSMs compared to their genomic representation by dividing the fraction of TFs or CSMs by the genomic fraction of TFs (6.8%) or CSMs (6.8%). We plotted the distribution of these enrichment values for various values of N (Figures 7H).

**Identification of unique marker genes for PN subtypes in Figure 7.** We sought to identify unique markers for each *GH146*+ PN subtype. We formed populations that each consist of *GH146*+ cells belonging to a cluster identified using ICIM and t-SNE (Figure 2D). We then performed differential expression analysis comparing each cluster to all other *GH146*+ cells. We selected genes that were differentially expressed at a significance level of P < 0.05 after the Bonferroni adjustment for multiple testing and having median expression within the cluster of interest of >7 CPM, resulting in 1103 genes. We then filtered for genes that were identified as significantly enriched in only one cluster, resulting in 257 genes. This step was necessary because some genes were identified as significantly enriched in multiple clusters, which is consistent with reuse of genes as identity factors within a combinatorial code. Finally, we identified genes that were robust and unique markers for a single cluster. To do this, we calculated the fraction of cells within each cluster expressing a given gene at >7 CPM. We then filtered for genes that were expressed in >50% of the cells within a given cluster and in <10% of the cells in any other cluster. We plotted the distribution of expression levels of these genes in each cluster (Figure S6A). We also tried different ways to relax the criteria. For example, Figure S6B shows the result when we required that the gene is expressed in >50% of the cells (>7 CPM) within a given cluster and in <25% of the cells in any other cluster.

Technical artifacts such as dropouts can hinder the identification of unique markers. We therefore estimated the probability that our failure to identify unique markers can be accounted for by dropout. To do so, we assumed that each of the 30 molecularly distinct *GH146*+ PN subtypes expresses a single unique marker gene at a low level of expression (7 CPM). In our data, genes expressed at an average level of 7 CPM are not detected in ~60% of cells. We may fail to detect a gene because (1) the cell is not expressing the gene or (2) because of noise in gene expression or technical dropout. We therefore can place an upper bound on the probability of dropout of a gene that is expressed at 7 CPM at 60%. Our method for identifying unique markers requires that the gene is detected in 80% of cells within a cluster. The probability of failure to detect the marker gene for a given cluster is given by the probability of dropouts in 20% of the cells in a cluster. This probability is:

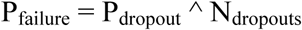

where P_dropou_t is estimated to be 60% and N_dropouts_ = 0.2 * N_cells_ is the number of cells in which the gene must drop out. We calculated this probability based on the number of cells Ncells in every cluster, ranging from 5 to 108. Then we calculated the probability that 25 out of 30 clusters do not have a unique marker gene by multiplying the probabilities of failure in 25 randomly sampled clusters. We performed this sampling 10,000 times and report the average (10^−62^). This value represents the probability of failing to detect a unique marker gene for 25 out of 30 clusters given that each cluster is expressing a single unique marker at 7 CPM.

We note that these calculations are conservative in several regards. First, we assumed that each cluster expresses a single marker gene. Realistically, each subtype may express multiple unique markers. This would increase the probability of detecting at least one of them. Second, unique markers may be expressed at levels higher than 7 CPM. We observed that the unique marker genes that we discovered are expressed at levels well above 7 CPM and biologically it is unlikely that a type identity factor would be expressed at extremely low levels. Thus, we estimate that our failure to detect marker genes can be explained by dropouts alone is very small.

**Information theory-based analyses of cell type identity related to Figure 7.** We sought to identify a minimal sets of genes that can encode the subtype identity of *GH146*+ PNs in a combinatorial fashion. Our motivation was to determine by direct search whether such a molecular combinatorial code exists. We also sought to characterize the redundancy of minimal combinatorial codes and the redundancy of individual genes in the context of such minimal codes.

To address this, we devised an algorithm that finds a minimal set of genes that is sufficient to encode the subtype identity of cells in a combinatorial manner based on ideas from information theory. An introduction to information theory is outside the scope of this work. Nevertheless, we provide a brief description of the basic concepts. Then we describe the algorithm and how it was applied in this work.

Entropy measures the uncertainty of a random variable (Shannon, 1948, Cover and Thomas 2006). Conditional entropy measures the uncertainty of a random variable given the knowledge of another variable (i.e., after conditioning on another variable). Conditioning on data never increases uncertainty (on average), which agrees with our intuition that additional information never hurts.

We use the notion of entropy H(C) to describe the uncertainty of cell type classification C. We use conditional entropy to describe the reduced uncertainty in classification due to knowledge of the expression state of a gene, H(C|G). The information gain due to knowledge of the expression state of the gene is the mutual information between the gene and the classification I(G;C), which can be defined as

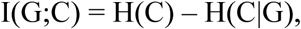

where H(C) is the entropy of cell type classification (without knowledge of the expression states of any genes) and H(C|G) is the entropy of cell type classification after conditioning on the expression state of gene G. Mutual information I(G;C) describes how much our uncertainty about classification C decreases when we observe the gene G. Mutual information I(G;C) can also be calculated directly from the probability distributions of cell type classes and expression states. For two discrete random variables G and C with their joint probability density function (pdf) p(x,y), the mutual information of G and C is defined as

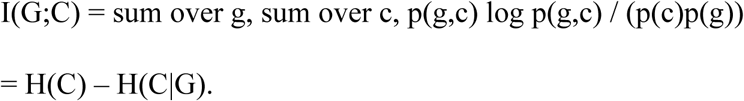

We often calculate the information content of a gene G with respect to the cell type classification C using this equation. Throughout this work, the base of the logarithm is 2 and so the unit of entropy and information is bit.

Information can be redundant. When we observe the expression state of a gene G, it may reduce the uncertainty of classification by an amount up to the information of that gene I(G;C). However, if we have already observed the expression states of other genes G’, then the additional observation of gene G may reduce the uncertainty of classification by an amount less than I(G;C),

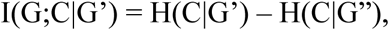

where G” is the set of genes containing G and G’. This is consistent with our intuition that the information carried by a gene G may be redundant with information that is already carried by genes G’. We define the redundancy R of a gene G in the context of a set of genes G’

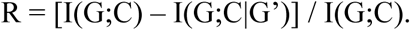

This describes the fraction of the information carried by the gene alone that it actually contributes when it is added to an existing set of genes (i.e., an existing code). R typically ranges from 0 to 1, where R = 0 means no redundancy and R = 1 means complete redundancy. In some special cases, there is synergy in the multivariate mutual information which causes R < 0.

We now describe the algorithm for finding a minimal combinatorial code. This problem is closely related to the Feature Reduction n-k (FRn-k) problem in machine learning (Battiti 1994). To solve this problem, we employ a greedy algorithm using mutual information similar to that described in (Kwak and Choi, 2002). The problem is formulated as follows:

Given an initial set F with n features and set C of all output classes, find the subset S ⊆ F with k features that minimizes H(C|S), which is equivalent to maximizing the mutual information I(C;S).

Our algorithm is as follows:

1. Initialize S as the empty set and F as the initial set of n features.
2. For all f_i_ in F, compute I(C;f_i_).
3. Find the feature f_i_ that maximizes I(C; f_i_). Add f_i_ to set S. Remove f_i_ from set F.
4. Repeat until the desired number of features k is selected:

1. For all f_i_ in F, compute I(C;f_i_|S).
2. Find the feature f_i_ that maximizes I(C; f_i_|S). Add f_i_ to set S. Remove f_i_ from set F.
5. Return the set S containing the selected features.

We repeat this computation with increasing k until the output set S explains a chosen amount of uncertainty in the classification C. Typically, we choose this termination condition as 99% of the entropy H(C) of classification C.

We note that the computation of mutual information is dramatically more efficient when G and C are discrete. We therefore binarized expression levels using a cutoff of log_2_(CPM+1) =3. This cutoff was chosen based on the minimum in the distribution of expression levels across all genes and all cells (Figure S7A). We varied the cutoff value between 2 and 6 and found that our results were essentially unchanged. The compactness of the minimal codes for *GH146*+ PN subtype identity and the genes included in the code were nearly identical to that obtained using the cutoff of 3 (data not shown). We also calculated the correlation between the information carried by each gene under different values of the cutoff with the information carried under the cutoff of 3 (Figure S7B). This analysis revealed that the information content of genes is not very sensitive to the precise choice of cutoff for binarization across the range of 2 to 6. We also found that other discretization schemes, such as a different number of levels of expression (e.g. Off, Low, Medium, High), yielded similar results (data not shown). All calculations related to information theoretic analyses were performed using binarized expression levels with a cutoff of log_2_(CPM+1) = 3.

**Application of information theory-based approaches to combinatorial coding of PN subtype identity in Figure 7.** We initially applied the algorithm to *Mz19*+ PN cells to test whether it is capable of identifying a set of genes that is sufficient for a combinatorial code of cell type identity. We formed a population consisting of the 175 *GH146*+ cells that belong to the classes labeled by *Mz19-GAL4* (108 DA1, 35 VA1d, and 32 DC3 cells). We created a binary expression matrix consisting of the ON/OFF states of all 15,522 genes that remained after removing genes that were not detected in any cells. We calculated the mutual information of each gene with respect to the *GH146*+ subtype classification. We then used the greedy algorithm described above to find a minimal set of genes for encoding *GH146*+ subtype identity (Figure 7A). The initial set of features F was the top 30 most informative genes among the 15,522 genes in the expression matrix.

We next applied this approach to all *GH146*+ PNs. We formed a population consisting of the 902 *GH146*+ cells belonging to the adPN and lPN lineages (Figure 2D). We created a binary expression matrix consisting of the ON/OFF states of all 15,522 genes that remain after removing genes that are not detected in any cells. We calculated the mutual information of each gene with respect to the *GH146*+ subtype classification (Figure 2D). We used the greedy algorithm described above to find minimal sets of genes for encoding *GH146*+ subtype identity with k varying from 1 to 20. The initial set of features F was the top 30 most informative genes among all 15,522 genes (genome-wide), among the 1062 TFs, or among the 1048 CSMs. We plotted the uncertainty explained by the minimal codes obtained with each value of k (Figure 7B). We then chose minimal codes that explained 95% of the uncertainty of *GH146*+ subtype classification (Figures 7C–E).

To evaluate whether TFs and CSMs carry more information than other genes, we found minimal sets of genes using an initial set of features F consisting of 1,000 genes chosen at random from among the 13,608 expressed in the genome after excluding the genes annotated as TFs and CSMs. We performed this search with 100 replicates. We plotted the mean uncertainty explained at various values of k and the standard deviation across the replicates (Figure 7B).

We assessed the redundancy of individual genes in the context of the minimal code. To do this, we calculated the redundancy R when each gene was added to the minimal code identified from all genes (genome-wide). The redundancy R is defined as the fraction of the information carried by the gene alone I(G;C) that it actually contributes when it is added to the minimal code I(G;C|G’),

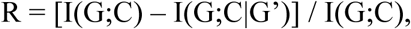

where G’ is the set of genes in the minimal code. R typically ranges from 0 to 1, where R = 0 indicates no redundancy and R = 1 indicates complete redundancy. However, in rare cases, genes had negative redundancy, indicating synergy in the multivariate mutual information. All of these genes had low information (I(G;C) < 0.05 bits). We excluded these genes from the visualization (Figure 7F). We calculated the fraction of highly informative and redundant genes that were annotated as TFs or CSMs.

## Acknowledgments

We thank Liming Tan and Larry Zipursky for sharing detailed protocols of cell dissociation; Gerry Rubin, Gerald Campbell, Adrian Moore, Ben White, Bloomington and Kyoto Stock Centers for reagents; Spyros Darmanis for discussions; Tom Clandinin, Jan Lui, Dan Pederick, Kang Shen, and Andrew Shuster for comments on the manuscript. H.L. is a Stanford Neuroscience Institute Interdisciplinary Postdoctoral Scholar, F.H. acknowledges support from the National Science Foundation Graduate Research Fellowship, S.R.Q. is a Chan Zuckerberg Investigator, and L.L. is an HHMI Investigator. This work was supported by NIH grant R01-DC005982 (to L.L.).

---

**Figure S1.**
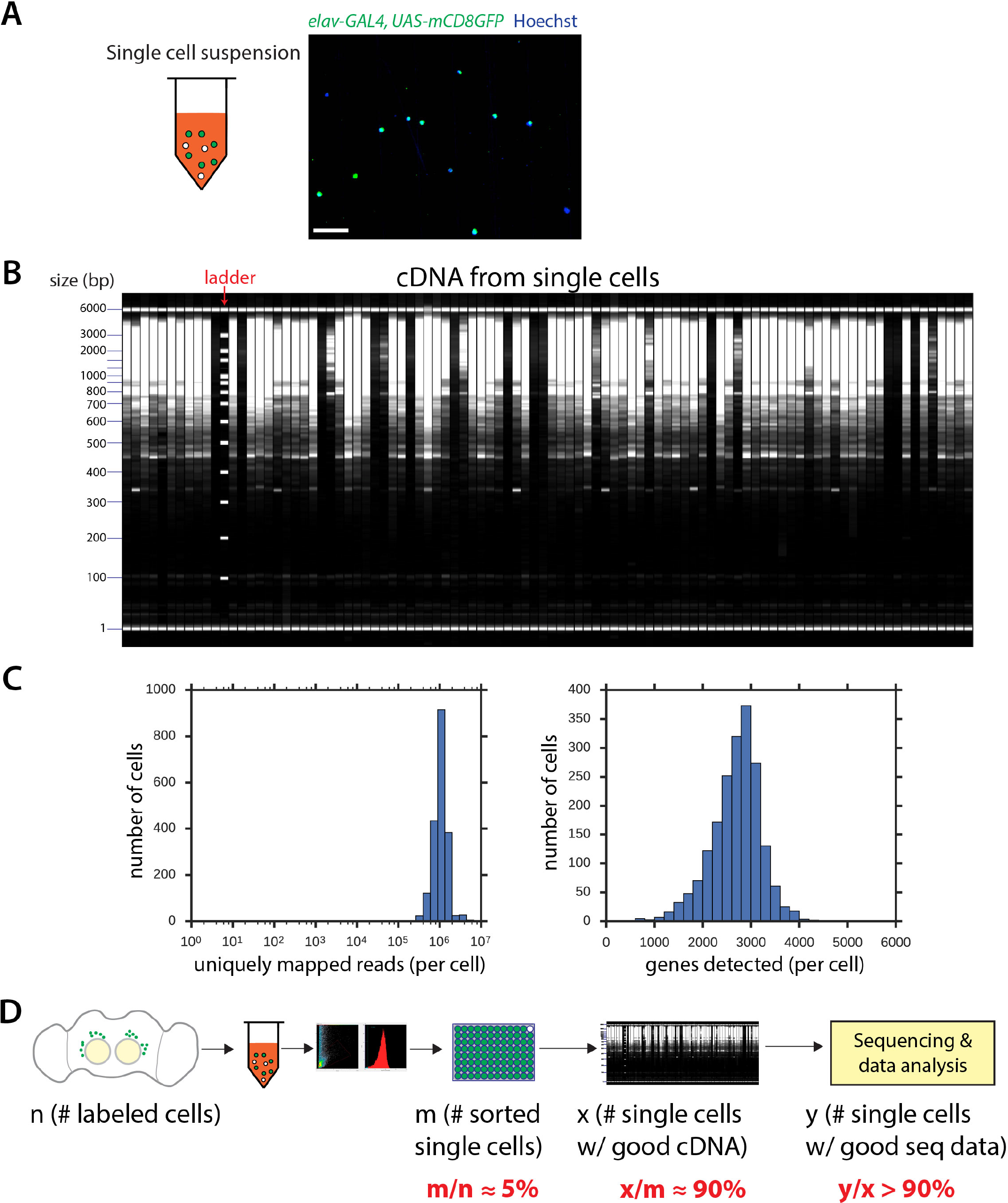
Single-cell RNA-seq Protocol for the *Drosophila* Pupal Brain, related to Figure 1. (A) Image of single-cell suspension after brain dissociation. Pupal brains were dissected and dissociated. Sample was imaged using epifluorescence microscopy. DNA was stained using Hoescht 33342 (blue). Scale bar, 50 µm. (B) Representative image of cDNA size distribution for 96 wells as measured using the Fragment Analyzer automated capillary electrophoresis system (Advanced Analytical). (C) Distributions of the number of uniquely mapped reads (left) and genes detected (right) per cell. On average, 1 million reads mapped uniquely to the *Drosophila* genome per cell, and 3000 genes were detected (CPM > 3). (D) Summary of efficiency for key steps of the single-cell RNA-seq protocol based on PNs. The efficiency of obtaining high-quality cDNA (x/m) from astrocytes is lower (~30%).

**Figure S2.**
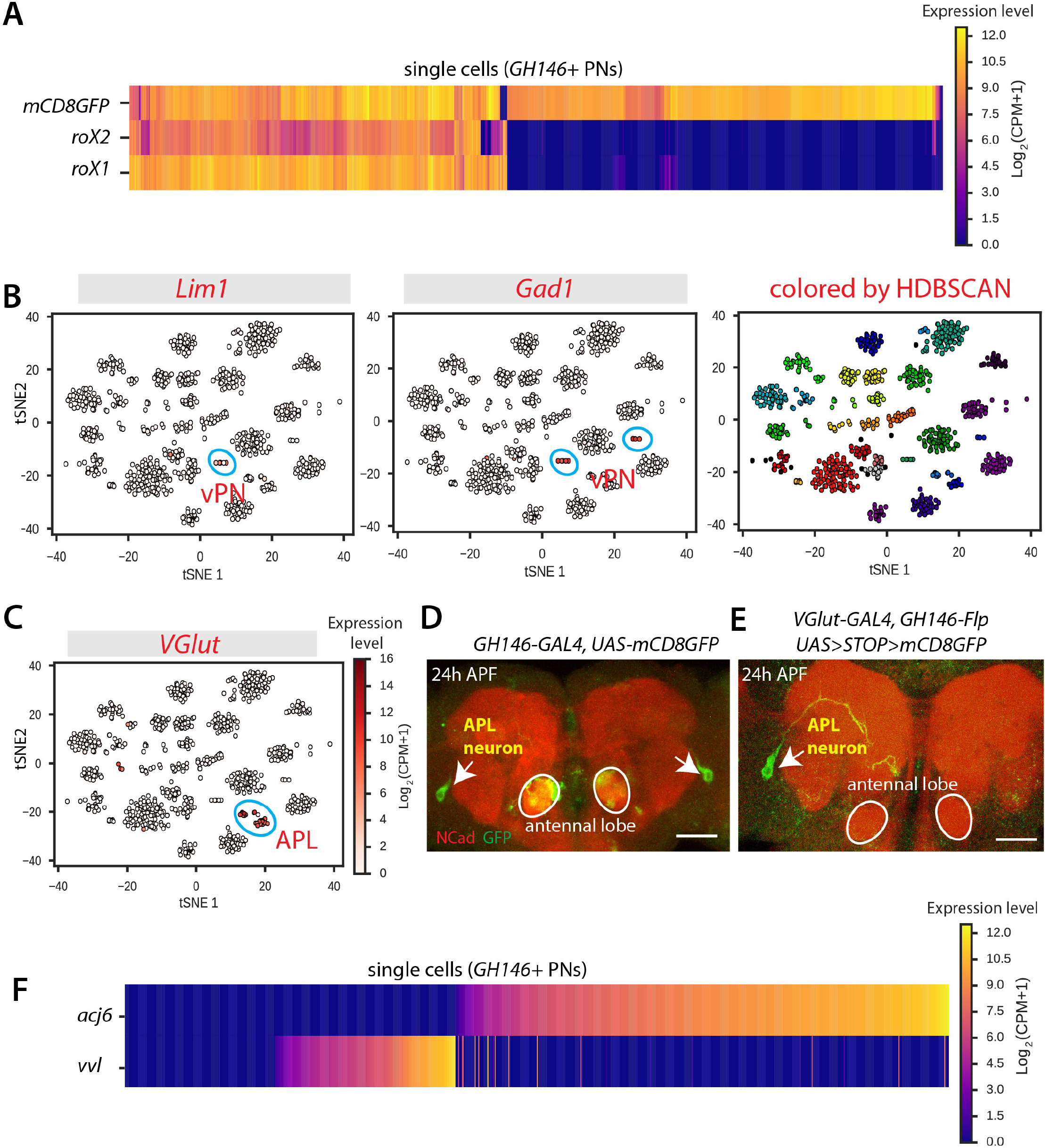
Single-cell RNA-seq Analysis for GH146+ Projection Neurons, related to Figure 2. (A) Heat map showing expression of the male-specific genes *roX1* and *roX2* in individual cells. Each column is one cell. As expected, ~50% of *GH146-GAL4*+ cells express these genes. Cells were ordered using hierarchical clustering. (B) Visualization of *GH146*+ PN cells using tSNE based on 158 genes identified using ICIM. Each dot is one cell. Cells are arranged according to similarity of expression profiles of the selected genes. Cells are colored by expression levels of *Lim1* (left), *Gad1* (middle) (see C for color bar; CPM, counts per million), and by HDBSCAN, which is a hierarchical density-based clustering algorithm (right). Two distinct clusters express *Gad1*, one of which expresses *Lim1*; both are unique to vPNs, indicating that these clusters correspond to *GH146*+ vPNs. (C) Visualization of *GH146*+ PN cells as in Figure S2B with cells colored according to expression of *VGlut* (CPM, counts per million). One distinct cluster expresses *VGlut* (outlined), which is a unique marker for anterior paired lateral (APL) neurons, indicating that this cluster contains APL neurons. (D) Confocal images showing that APL neurons (indicated by arrows) are labeled by *GH146-GAL4* driven *UAS-mCD8GFP*. Scale bar, 50 µm. (E) Confocal image showing that APL neurons (arrow) are labeled by *VGlut-GAL4* (after intersecting with *GH146-Flp)*. Ncad staining labels neuropil (red), and antennal lobes are outlined. (F) Heat map showing expression of the lineage-specific transcription factors *acj6* and *vvl* in *GH146*+ adPN and lPN cells (after removal of vPNs and APL neurons) (CPM, counts per million). Cells are ordered by *acj6* expression, and then by *vvl* expression.

**Figure S3.**
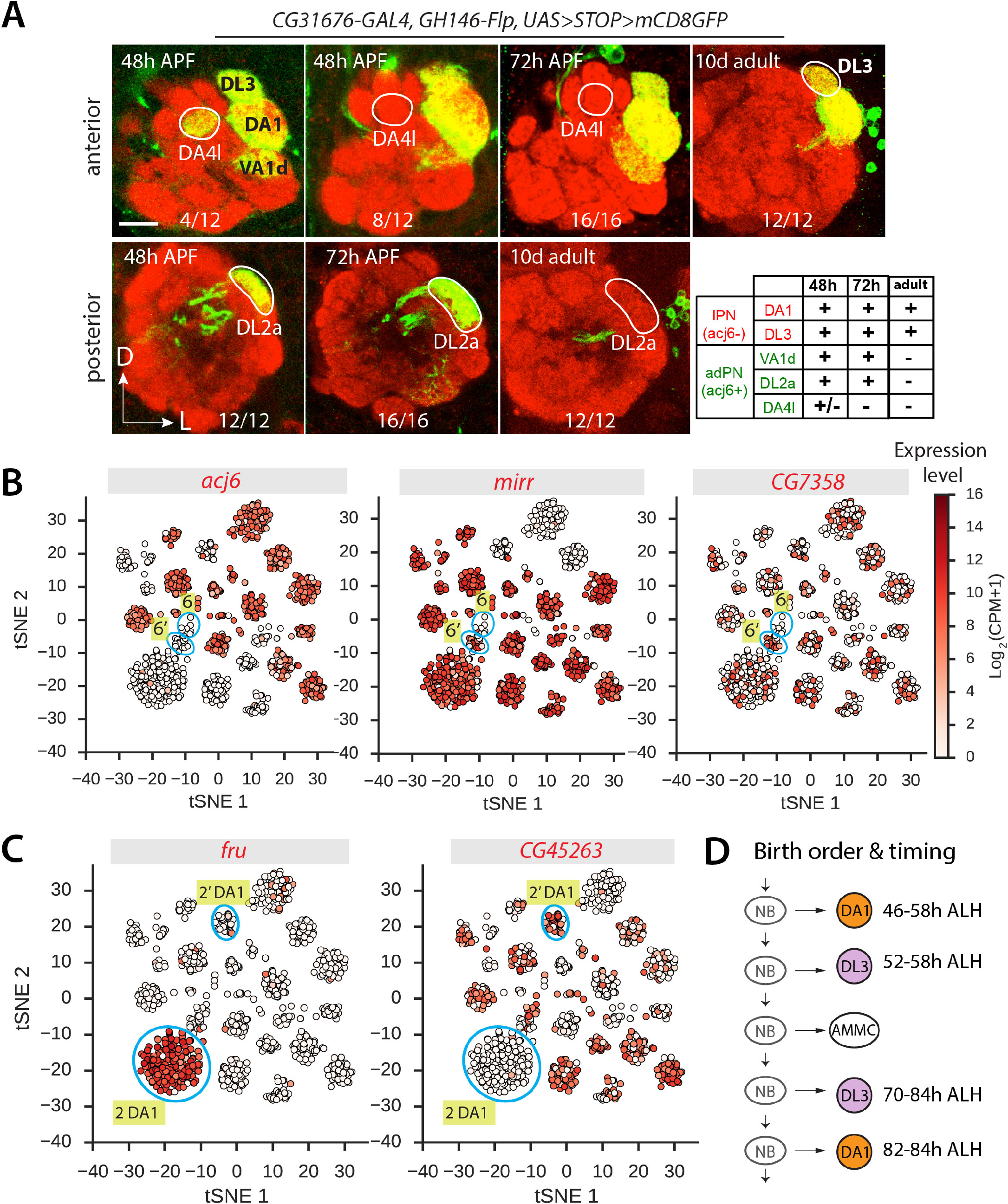
Assign Clusters to PN Classes Using Newly Identified Markers, related to Figure 4. (A) Systematic characterization of *CG31676-GAL4* expression in PNs after intersecting with *GH146-Flp* at 48h and 72h APF, as well as 10d adult. Expression patterns are summarized in the table: +, expressed; −, not expressed; +/−, expressed in a subset of flies. *CG31676-GAL4* stably labels DA1 and DL3 from pupa to adult. Note that *CG31676-GAL4* also transiently labels DL2a and DA4l adPNs (as summarized in the table), but we could not unambiguously map them to corresponding clusters. N-cadherin (Ncad) staining was used to label neuropil (red). Scale bar, 20 µm. (B) Visualization of *GH146*+ PN cells using tSNE as in Figure 4A showing expression levels of *acj6, mirr*, and *CG7358* (CPM, counts per million). Clusters #6 and #6’ are both *acj6−*. Cluster #6’, but not Cluster #6, is *mirr*+ and *CG7358*+. (C) Visualization of *GH146*+ PN cells using tSNE as in Figure 4A showing expression levels of *fru* and *CG45263* (CPM, counts per million; see color bar in Figure S3B). *fru* is expressed in Cluster #2, but not Cluster #2’, while *CG45263* is expressed in Cluster #2’, but not Cluster #2. Both Clusters #2 and #2’ map to DA1 PNs. (D) Schematic summary of birth order and timing of the lateral neuroblast (NB) lineage. Both DA1 and DL3 PNs are born in two different periods, separated by antennal mechanosensory and motor center (AMMC) neurons (Lin et al., 2012). ALH, after larval hatching.

**Figure S4.**
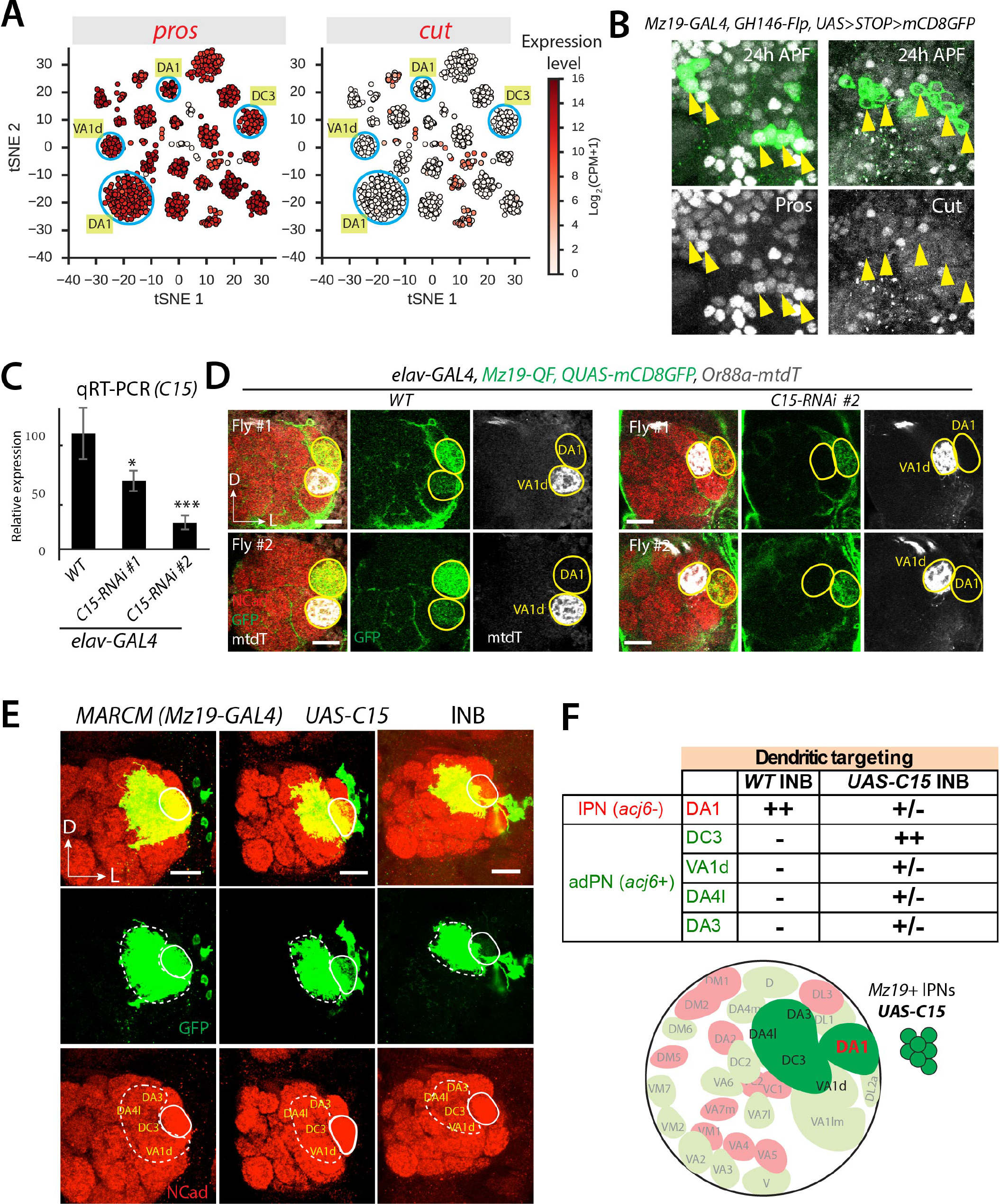
Validation of Transcription Factor Expression Patterns, related to Figure 5. (A) Visualization of *GH146*+ PN cells using tSNE as in Figure 4A showing expression of *prospero (pros)* and *cut* (*ct*). *pros* is expressed in *Mz19*+ PNs, while *ct* is not. (B) Antibody staining shows that *Mz19*+ PNs (arrowheads) express Pros but not Cut at 24 APF, consistent with RNA-seq data, as shown in (A). (C) Quantitative PCR (qPCR) measurement of the knockdown efficiency of two *UAS-C15-RNAi* lines. *elav-GAL4* was crossed with either *W^1118^* (control) or two *C15-RNAi* lines, and mRNA was extracted from 5-day-old adult fly heads (N = 3 x 10 heads for each condition). Expression levels are normalized to *actin5C*. Error bars show SEM. *, P < 0.05; ***, P < 0.001 (t test). (D) Two additional examples of dendrite targeting of *Mz19-QF*+ PNs in *WT* and *C15* knockdown, as in Figure 5C. DA1 and VA1d glomeruli are outlined in yellow. (E) Three additional examples of gain-of-function analysis of *C15* misexpression in *Mz19-GAL4*+ MARCM lateral neuroblast (NB) clones, as in Figure 5E. Mistargeted regions are outlined and corresponding glomeruli are indicated. (F) Summary of mistargeting phenotypes for *UAS-C15* lNB clones. ++, fully targeted; −, not targeted; +/−, partially targeted. Lower panel is a schematic of mistargeting phenotypes. Note that all mistargeted glomeruli are normally innervated by adPNs. N-cadherin (Ncad) staining was used to label neuropil (red). Scale bar, 20 µm.

**Figure S5.**
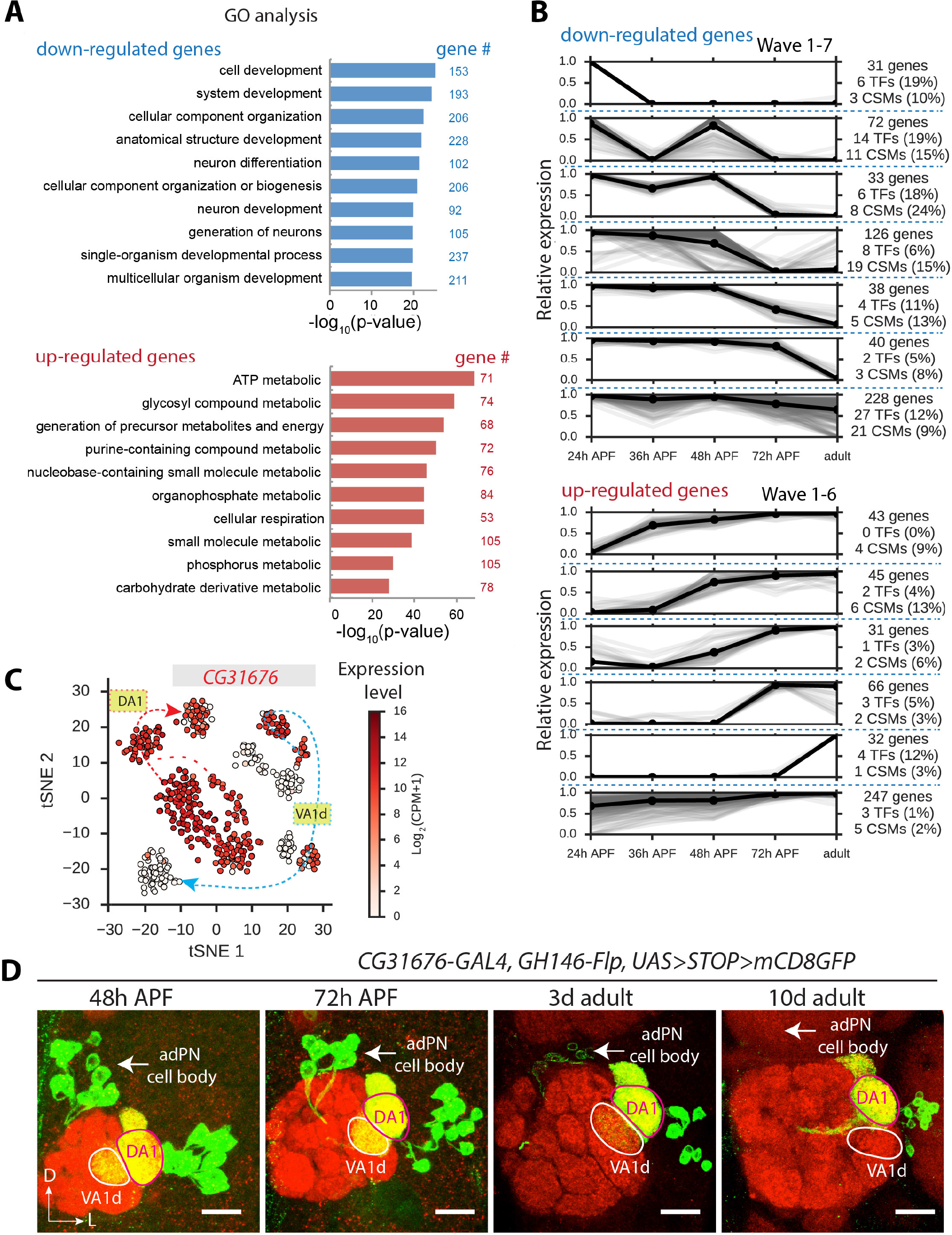
Analysis of *Mz19*+ PN Development and Maturation, related to Figure 6. (A) Gene Ontology (GO) analysis of genes that were upregulated and downregulated between 24h APF and adult in *Mz19*+ PNs (P < 10^−5^). For the top 10 most significantly enriched GO terms, the significance of enrichment and the number of genes corresponding to each term are shown. (B) Expression dynamics of the genes identified in (A) spanning the 24h, 36h, 48h, and 72h APF, and adult stages. Genes were classified based on their dynamical profiles (Experimental Procedures). This approach identified 7 distinct dynamical patterns of expression among the down-regulated genes (waves 1–7) and 6 such patterns among up-regulated genes (waves 1–6). For individual genes, the median expression level at each developmental stage is shown in gray (normalized to maximum expression across developmental stages). The mean expression profile across the genes assigned to a wave is shown in black. For each wave, the number and fraction of genes that were TFs and CSMs are indicated. (C) Visualization of *Mz19*+ PN cells from developmental stages ranging from 24h APF to adult as in Figure 6B showing expression of *CG31676* (CPM, counts per million). In DA1 PNs, *CG31676* is expressed at all stages. In VA1d PNs, *CG31676* is expressed at all pupal stages, but is turned off in adult. (D) Confocal images (anterior stacks) showing *CG31676-GAL4* expression after intersecting with *GH146-Flp*, at various pupal and adult stages. DA1 and VA1d glomeruli are outlined and adPN cell bodies are indicated (arrow). Consistent with our RNA-seq data (Figure S5C), *CG31676* is expressed at all time points in DA1 PNs, while in VA1d PNs *CG31676* is expressed at all pupal stages and then turned off in 10 day old adult flies. We observed very weak expression in 3-day-old adult flies, likely due to perdurance of mCD8GFP. Scale bar, 20 µm.

**Figure S6.**
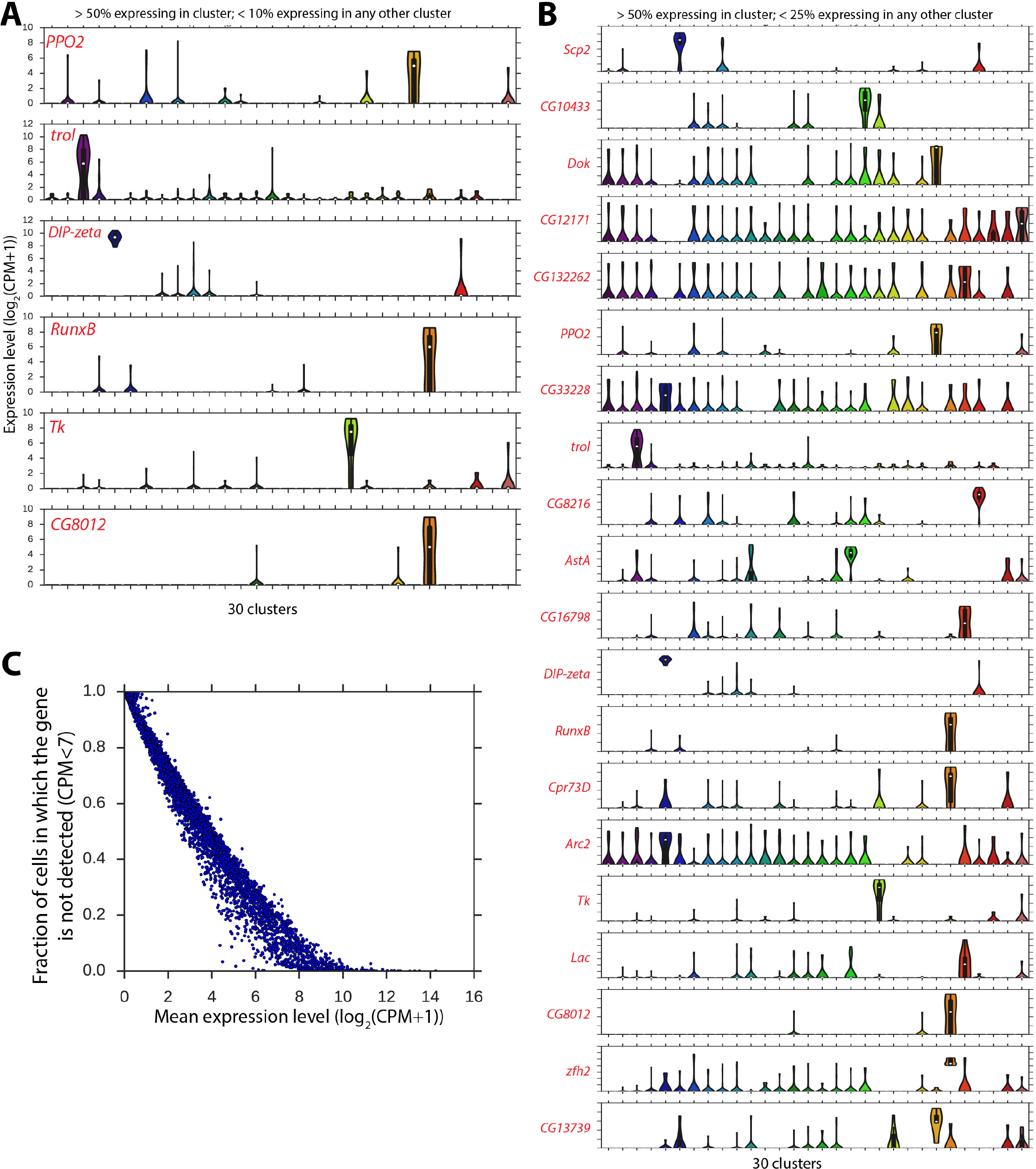
Searching for Unique Markers of Individual PN Subtypes, related to Figure 7. (A) Violin plots showing expression of genes identified as unique marker genes among cells within each *GH146*+ PN cluster. These 6 genes were identified using the criteria: (1) >50% cells within the cluster express the gene, and (2) <10% cells in any other cluster express the gene. Expression was defined as >7 CPM, or log_2_(CSM+1) > 3. These genes specify 5 distinct PN clusters. (B) Violin plots showing expression of genes that were identified as unique markers using less stringent criteria. The criteria used here were: (1) >50% of the cells in the cluster express the gene, and (2) <25% of the cells in any other cluster express the gene. Expression was defined as >7 CPM, or log_2_(CSM+1) > 3. 20 genes were found, but many of these genes are clearly not unique to a single cluster. This indicates that relaxing the stringency of our search results in many genes being discovered which are not in fact unique markers. See (A) for labels of the expression level. (C) Estimation of an upper bound on dropout rate in our single-cell RNA-seq measurement. Each dot is a gene. The mean expression level of the gene across all *GH146*+ PNs is plotted against the fraction of cells in which the gene is not detected (<7 CPM, or log_2_(CPM+1) < 3). Detection failure events can occur because either (1) the gene is not expressed in the cell, or (2) failure to detect expression of the gene despite the presence of mRNA transcripts due to technical artifact (called dropouts). Thus, the fraction of detection failure events offers an upper bound on dropout rate. We used this upper bound to calculate the probability that we failed to detect unique markers for 25 PN classes due to dropout alone (Experimental Procedures).

**Figure S7.**
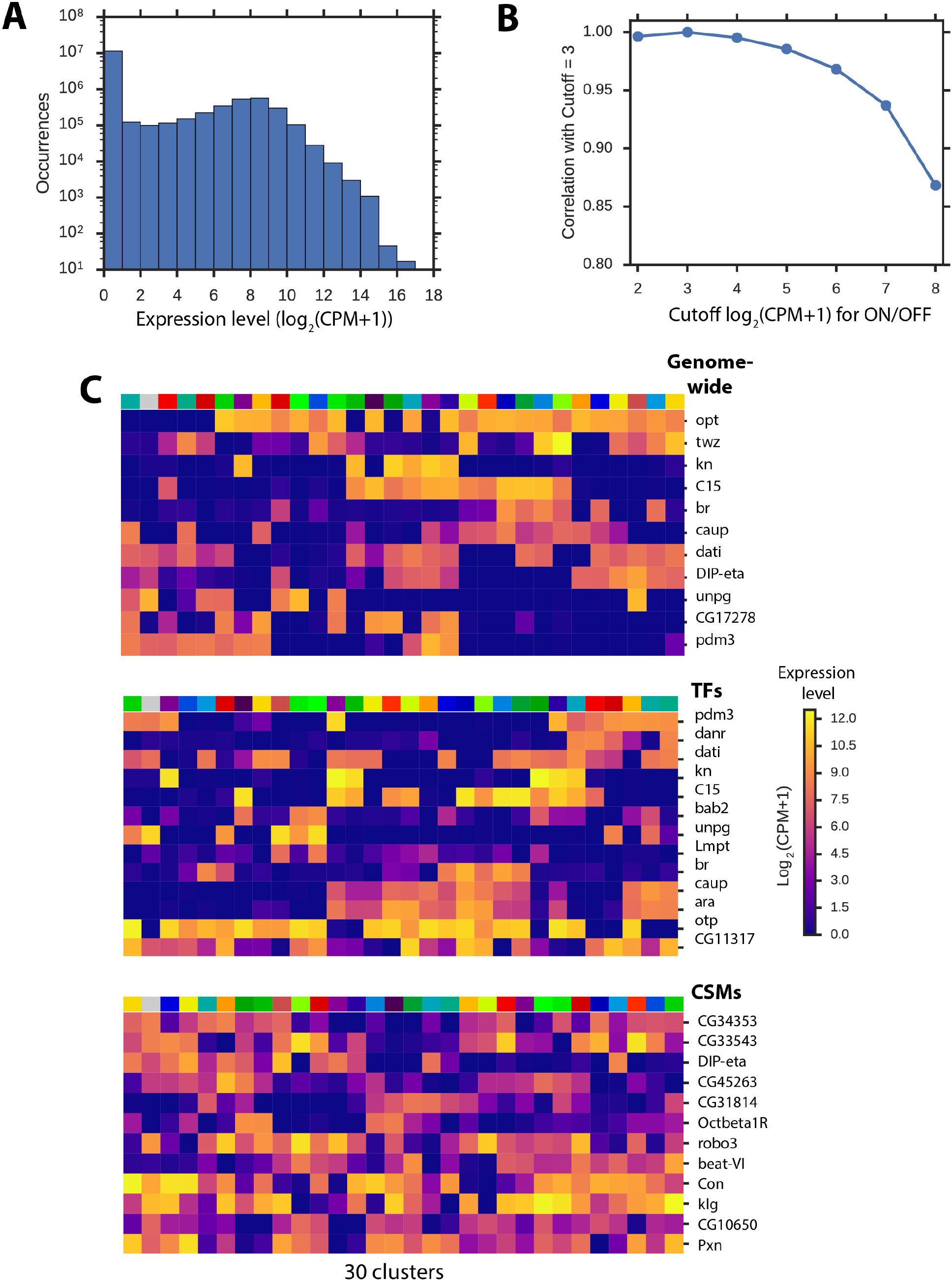
Combinatorial Molecular Codes of PN Subtype Identity, related to Figure 7. (A) Distribution of expression levels across all genes and all *GH146*+ cells (CPM, counts per million). We chose to binarize expression levels using a cutoff of log_2_(CPM+1) = 3 (equivalent to CPM = 7) because the distribution exhibits a distinct minimum at this value. (B) Robustness of information to the choice of binary cutoff. We calculated the Pearson correlation between the mutual information of a gene under various binarization cutoffs and the mutual information at the binarization cutoff that we chose (log_2_(CPM+1) = 3). For binarization cutoffs ranging from 2 to 6, the mutual information was highly similar (ρ > 0.95), indicating that the precise choice of binarization threshold does not affect our results. (C) Expression levels of genes in the minimal combinatorial codes for *GH146*+ PN subtype identity. Each column is a cluster and each row is a gene. Color indicates the median expression of the gene among the *GH146*+ cells within the cluster (CPM, counts per million). These plots correspond to Figures 7C-E before binarization. Cells and genes are arranged by hierarchical clustering on binarized expression states (dendrograms shown in Figures 7C-E).

